# AutoCryoPicker: An Unsupervised Learning Approach for Fully Automated Single Particle Picking in Cryo-EM Images

**DOI:** 10.1101/561928

**Authors:** Adil Al-Azzawi, Anes Ouadou, John J. Tanner, Jianlin Cheng

**Affiliations:** Electrical Engineering and Computer Science Department, University of Missouri, Columbia, MO 65211, USA; Departments of Biochemistry and Chemistry, University of Missouri, Columbia, MO 65211-2060, USA.; Informatics Institute, University of Missouri, Columbia, MO 65211, USA

**Keywords:** Clustering, Intensity Based Clustering (IBC), micrograph, Cryo-EM, singe particle pickling, protein structure determination

## Abstract

**Background:** An important task of macromolecular structure determination by cryo-electron microscopy (cryo-EM) is the identification of single particles in micrographs (particle picking). Currently, particle picking is laborious, time consuming, and potentially biased due to the need of human intervention to initialize the particle picking. The results typically include many false positives and negatives. Adjusting the parameters to eliminate false positives often excludes true particles in certain orientations. The supervised machine learning (e.g. deep learning) methods for particle picking often need a large training dataset, which requires extensive manual annotation. Other reference-dependent methods rely on low-resolution templates for particle detection, matching and picking, and therefore, are not fully automated. These issues motivate us to develop a fully automated, unbiased framework for particle picking.

**Results:** We design a fully automated, unsupervised approach for single particle picking in cryo-EM micrographs. Our approach consists of three stages: image preprocessing, particle clustering, and particle picking. The image preprocessing is based on image averaging, normalization, cryo-EM image contrast enhancement correction (CEC), histogram equalization, restoration, adaptive histogram equalization, guided image filtering, and morphological operations significantly improves the quality of original cryo-EM images. Our particle clustering method is based on an intensity distribution model which is much faster and more accurate than traditional K-means and Fuzzy C-Means (FCM) algorithms for single particle clustering. Our particle picking method, based on image cleaning and shape detection with a modified Circular Hough Transform algorithm, effectively detects the shape and the center of each particle and creates a bounding box encapsulating the particles.

**Conclusions:** AutoCryoPicker can automatically and effectively recognizes particle-like objects from in noisy cryo-EM micrographs without the need of labeled training data and human intervention and therefore is a useful tool for cryo-EM protein structure determination.

## Background

For decades, X-ray crystallography has been the dominant technique for obtaining high-resolution structures of macromolecules. Single-particle cryo-electron microscopy (cryo-EM) was traditionally used to provide low resolution structural information on large protein complexes that resisted crystallization (e.g., highly symmetric particles of viruses). Though the basic workflow of cryo-EM has not changed considerably over the years, recent technological advances in sample preparation, computation and especially instrumentation have revolutionized the field of structural biology [1] [2] [3], allowing it to solve large protein structures at better than 3 Å resolution [4] [5] [6] [7].

Cryo-EM micrographs contains two-dimensional projections of the particle in different orientations. Generally, cryo-EM images have low contrast, due to the similarity of the electron density of the protein to that of the surrounding solution, as well as the limited electron dose used in data collection. In addition, the micrographs may contain sections of ice, deformed particles, protein aggregates, etc., which can complicate particle picking. Because a large number of single-particle images, extracted from cryo-EM micrographs are required for a reliable 3D reconstruction of the underlying structure, particle recognition thus, represents a significant bottleneck in cryo-EM structure determination.

To address the bottleneck, numerous computational approaches have been proposed to facilitate the particle picking process [8] [9] [10] [11] [12] [13] [14]. These methods can roughly be divided into two categories: generative methods [15] [16] [17] and discriminative classification methods [18] [19] [20] (e.g. the recent deep learning methods [21] [22]). The generative methods measure the similarity of an image region to a reference to identify particle candidates from micrographs. A typical generative method employs a template-matching technique with a cross-correlation similarity measure to accomplish particle selection. The discriminative methods first train a classifier on a labeled dataset of positive and negative particle examples, then apply it to detecting particle images from micrographs images.

DeepPicker [21] is a deep learning method for semi-automated particle selection and picking. The first part of the method involved the manual creation of training data. The second part was fully automated by learning patterns from the training data to classify particles. DeepEM [22] uses a convolutional neural network (CNN) to recognize particles. The CNN was trained on a manually curated dataset. The training dataset was augmented by adding additional particles images generated by image rotation.

The existing unsupervised approaches distinguish the particle-like objects from background noise in micrographs via an unsupervised learning manner without the need of any labeled training data [10] [11] but, they do not fully exploit the intrinsic and unique characteristics of particles to facilitate automated particle picking. Therefore, the unsupervised approaches are often combined with the reference template matching or classification-based approaches to achieve good picking results. However, in this case, the training dataset has to be manually created to train the model. Although these approaches have greatly reduced time and effort spent on single-particle data analysis, most of them are not fully automated and still require substantial human intervention to initialize the particle selection process. For instance, most methods require users to prepare an initial set of high-quality reference particles used as templates to search for similar particle candidates from micrographs, while the discriminative approaches usually demand the user to manually pick a number of positive and negative samples to train the classifier first.

In this paper, we develop a fully automated approach for particle picking (AutoCryoPicker) that is based on advanced image preprocessing, robust clustering based on the intensity distribution, and sophisticated shape detection. The experimental results demonstrate that the fully automated particle picking scheme can accurately detect a number of particles that is comparable to those picked manually. The clustering method is also more accurate than k-means and Fuzzy C-means (FCM) for particle clustering. Therefore, our new automated picking approach can significantly reduce time and labor spent on single-particle data analysis and thus greatly relieves a bottleneck in the automated cryo-EM structure determination pipeline.

## Methods

Our AutoCryoPicker framework for automated particle picking is shown in Figure 1. In this framework, a user is not required to manually pick any particle from the micrographs. The fully automated approach has three main stages: preprocessing, clustering, and particle picking. In the preprocessing stage, several image processing methods are applied to enhance the input cryo-EM images such as image normalization, Contrast Enhancement Correction (CEC), etc. Clustering is done using three different algorithms k-means [23], Fuzzy C-Means (FCM) [24], and a new robustness clustering algorithm that addresses some typical clustering issues such as cluster destabilization due to random initialization of cluster centers. In the particle picking stage, a final set of particles is selected from clustered particle candidates.

**Figure 1.**
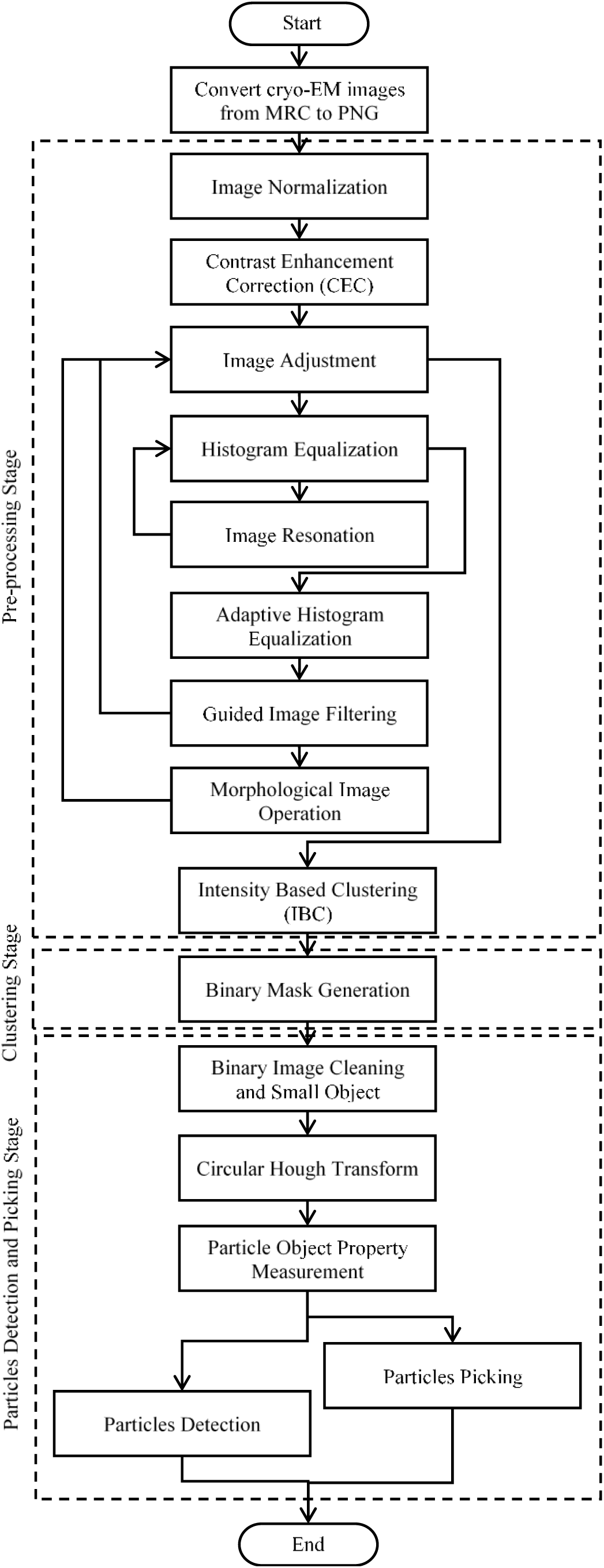
The general framework of AutoCryoPicker: Fully Automated Single Particle Picking. The dashed boxes represent three stages of the approach: pre-processing, particle clustering, and particle detection and picking. A solid box denotes an analysis step.

### Stage 1: Pre-processing

A standard cryo-EM image is stored in the MRC format, which defines a three-dimensional grid (array) of voxels each with a value corresponding to electron density or electric potential. In order to apply various image preprocessing techniques to improve the quality of noisy cryo-EM images, we convert cryo-EM images in the MRC format into widely used 16-bits PNG format using EMAN2 [25]. Since our goal is to use the unsupervised learning algorithm to cluster pixels based on the difference in intensity levels in any cryo-EM image, we select a set of advanced preprocessing tools to improve the quality of cryo-EM images. Those tools are tested on two different datasets.

There are two benefits of using the preprocessing. Firstly, those tools improve the contrast of the cryo-EM images by increasing the particle’s intensity. Secondly, pre-grouping the pixels inside each particle makes them easier to be isolated by the clustering algorithm. Specifically, the preprocessing tools are selected based on three main objectives: enhancing the global contrast of the cryo-EM, enhancing the local contrast and increasing the intensity level of each particle, and enhancing the particle shapes inside the cryo-EM images. In order to improve the entire contrast between particles and the background, image normalization is used first and then contrast enhancement and correction is applied to increase the global intensity value. To increase the global image contrast, histogram equalization is applied to enhance the pixel intensity level and then image restoration is used to recover and improve the quality of an image. To improve the local contrast and enhancing the definitions of edges in each particle, adaptive histogram equalization is employed. Moreover, guided image filtering is used to perform edge-preserving smoothing of each particle in the cryo-EM image. Finally, morphological image operation is applied to enhance the particle shape and make the particle regions similar to each other and different from the background regions. These preprocessing methods are described in details in the following steps.

### Step 1: Cryo-EM Image Resolution Improving

Cryo-EM images are affected by different factors that either corrupt the micrograph image signal by some gaussian noise or the image resolution. Different cryo-EM images have different artificial objects such as ice, which in some cases, have different thickness and similar ranges of the particle’s pixel intensity value. In this case, in a single cryo-EM image, a small number of particles may not have significant difference of scatter power. Technically, the cryo-EM image resolution can be improved using computational image (signal) averaging based on blur motion elimination. This is selected as a main step of the contrast transfer function (CTF) based on the image quality evolution of the single particle cryo-EM and 3D reconstruction tool of viruses [26].

Different cryo-EM images have different intensity value ranges. In order to unify the range values, we renormalize the micrograph by setting the background mean to zero and background variance to one. In this normalization, the pixel values become the Z-score, i.e., the number of sigma’s above noise level as shown in Equation (1) [27]:

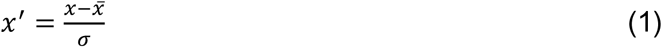

where 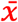 is the mean of the intensity pixel values, and *σ* is the standard deviation. For instance, for an image consisting of 50 frames, we used the image averaging and normalization function in EMAN2 [25] to average the 50 frames, resulting in a converted cryo-EM image for further processing and analysis as shown in Figure 2.

**Figure 2.**
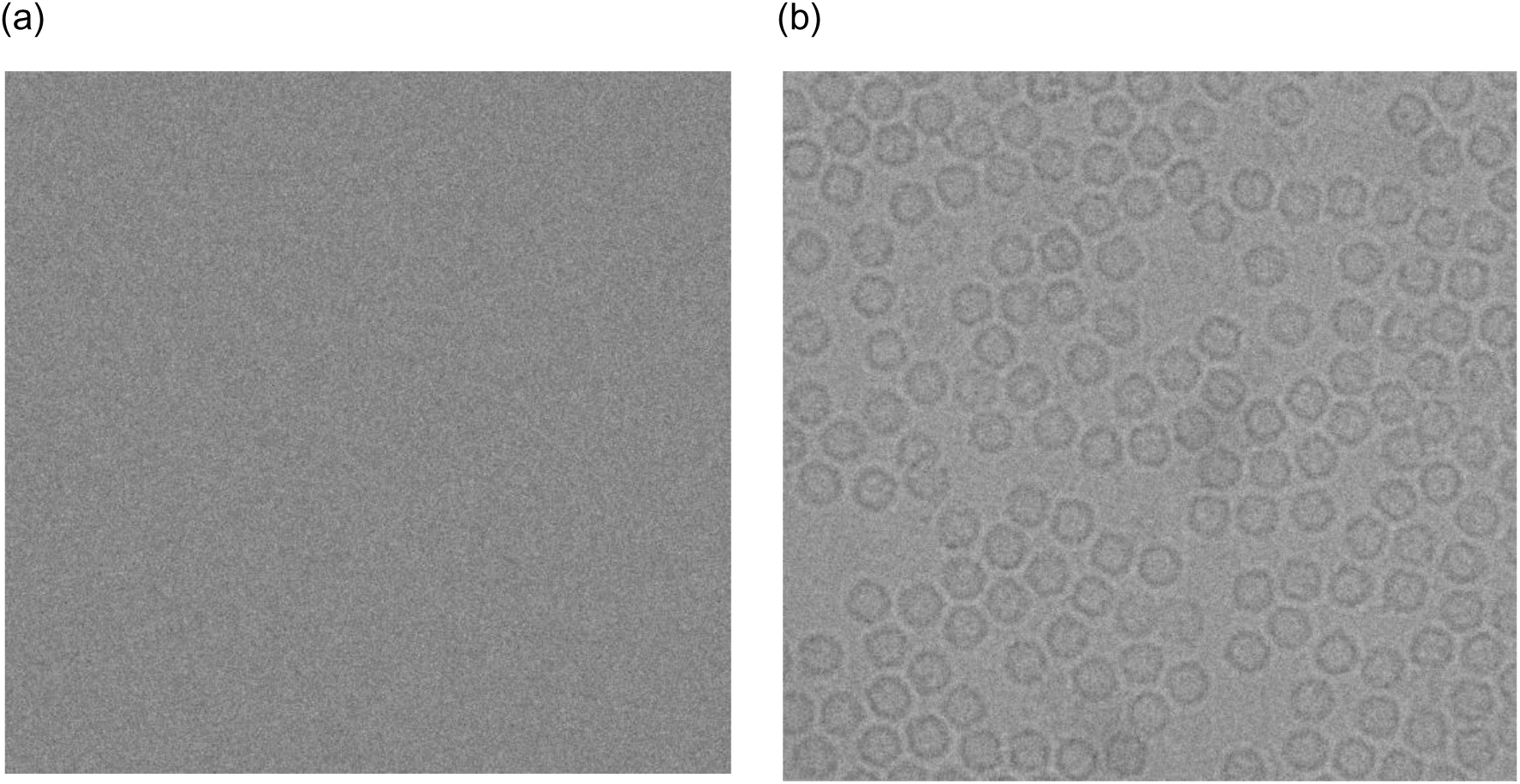
Cryo-EM image averaging and normalization result using EMAN2. (a) the original cryo-EM image (stack of 50 frame) in the MRC format before the averaging and normalization processing, (b) the cryo-EM image in PNG file format (single frame) after the averaging and normalization processing using EMAN2.

### Step 2: Global Cryo-EM Intensity Adjustment

Low-dose micrograph imaging models the exposure to a very low intensity beam in a large defocus area that has both good particle distribution and thin ice. This imaging mode produces very low intensity cryo-EM images. To overcome this problem, intensity adjustment is applied to map the cryo-EM image’s intensity values to a new range. An Intensity Enhancement Correction (IEC) procedure is used to identify the descent image intensity and improve signal to noise ratio in cryo-EM images. In order to enhance the global intensity adjustment, we apply three different steps.

1. Find Limits to Contrast Stretch: In this step, the range of image intensity is specified by detecting the low and high values via a MATLAB function “*stretchlim*”, which returns a two-element vector that consists of the low and upper intensity limits as shown in the cryo-EM histogram in Figure 3(a). By default, values in low and high intensities specify the bottom 2% and the top 2% of pixel values. In this case, the intensity level of each cryo-EM should be unified. The gray values returned can be used by the “*imadjust*” function [28] to increase the contrast of an image as shown in Figure 3(b).
2. Mid-Range Stretching: In this step, the cryo-EM image intensity values are stretched to improve their quality. The gray scale image pixels are mapped into the range [0 1] by dividing the intensity values of each pixel as shown in Equation (2).

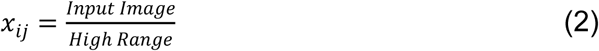

where *i* and *j* are the row and column index of cryo-EM image matrix respectively and the *High Range* is the highest intensity value in the input image. Figure 4(a) shows an original cryo-EM image, Figure 4(b) the histogram of the original image, Figure 4(c) a cryo-EM image after mid-range stretching and Figure 4(d) the histogram of the stretched image. The histogram in Figure 4(d) is more stretched than the original one in Figure 4(b).
3. Intensity Adjustment: The intensity values of the cryo-EM image are adjusted to new values in a condensed smaller range by using the MATLAB function “*imadjust*” [28]. Figure 4(e) shows an example of a cryo-EM image with contrast enhancement correction (CEC) and image adjustment, and Figure 4(f) shows the histogram of Figure 4(e) where the histogram looks more stretching and the contrast of the cryo-EM is enhanced compared with the original image in Figure 4(a).

**Figure 3.**
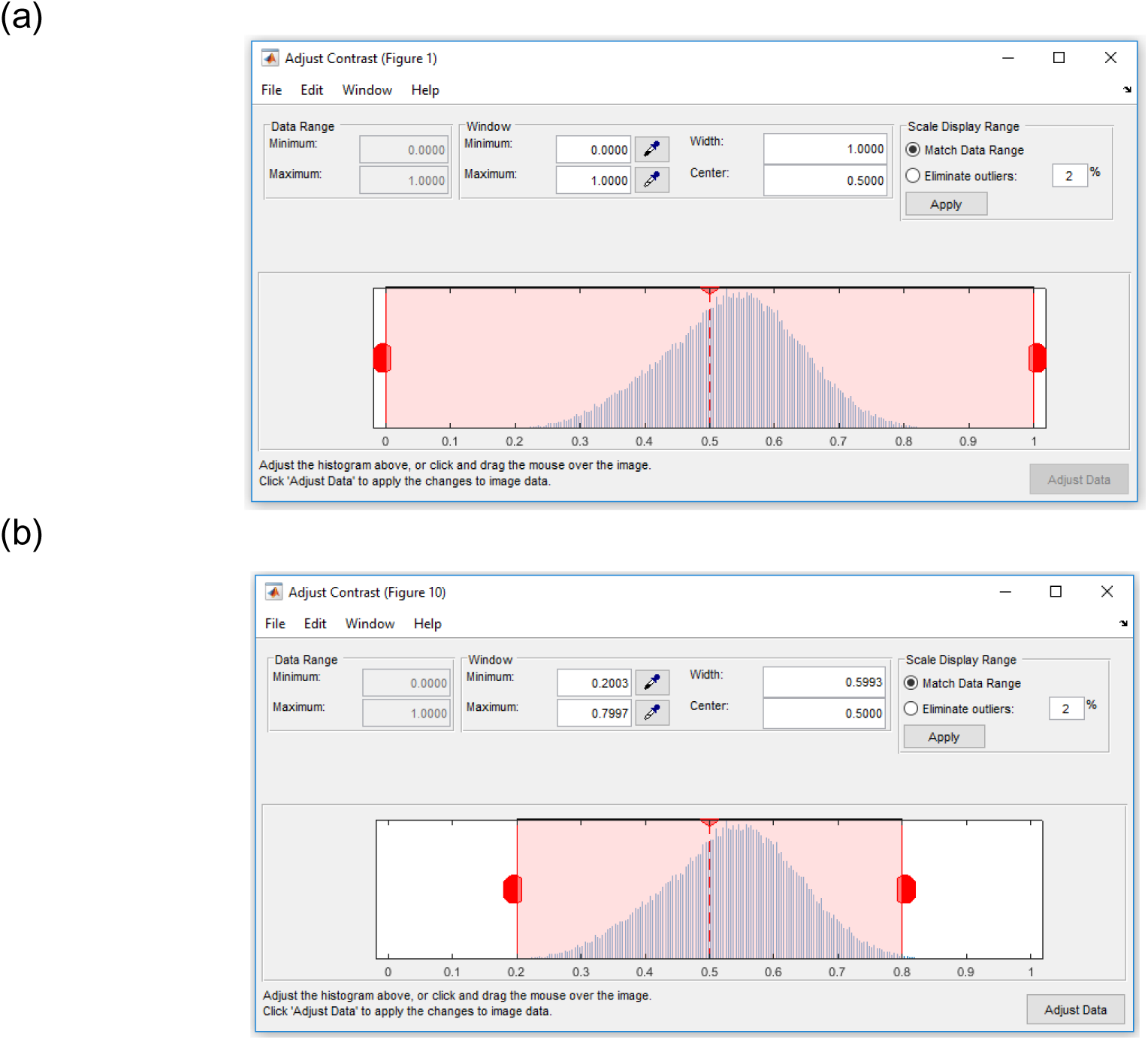
Contrast transfer correction and adjustment process. (a) illustration of the cryo-EM image histogram after the averaging and normalization step using EMAN2 and the a two-element vector that consists of the low and the upper intensity limits by default. The values in low_high specify the bottom 2% and the top 2% of all pixel values, (b) illustration of the cryo-EM histogram (Histogram shrinking) after automatically detecting and specifying the low and high intensity range (e.g. [0.2-0.8]).

**Figure 4.**
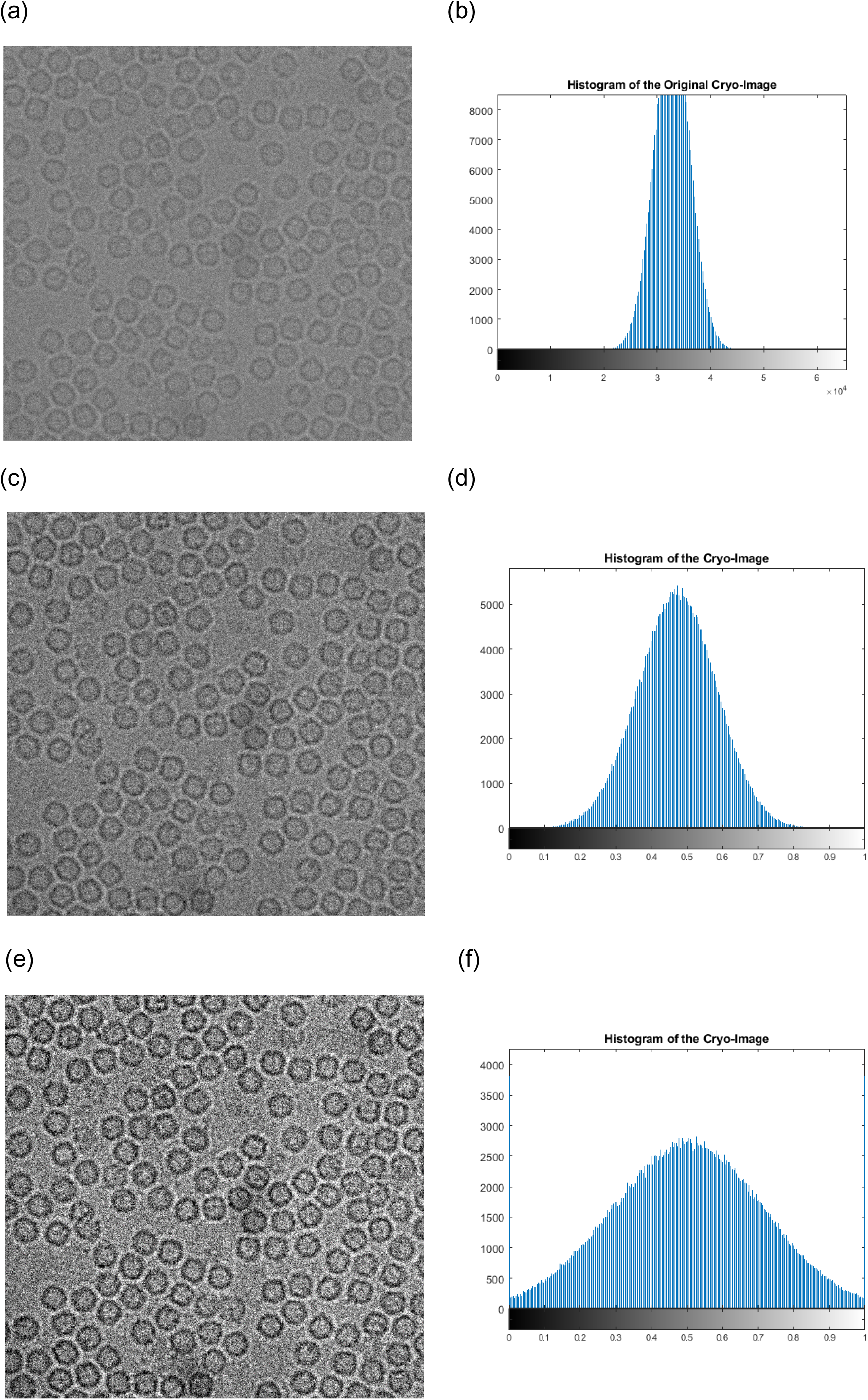
Cryo-EM Contrast Transfer Correction (CTC) process. (a) the original cryo-EM image after the applying the averaging and normalization process through the EMAN2 software, (b) Histogram of the original cryo-EM image, (c) The cryo-EM image after applying the mid-range stretching based on the low-high intensity range, (d) Histogram of the image in (c), (e) The cryo-EM image after applying the contrast enhancement correction (CEC) and image adjustment, (f) The histogram of the cryo-EM image after applying the contrast enhancement correction (CEC).

For better demonstrating the effects of the preprocessing steps, we zoom-in one particle image from different datasets. Figure 5(a) and (i) show two original particle images from two different datasets. Figure 5(b) and (j) show the cryo-EM Image resolution being improved by image averaging and normalization. We can notice that image noise has been reduced. Figure 5(c) and (k) illustrates the same single particle images after the global intensity adjustment using Intensity Enhancement Correction (IEC). In comparison with the same particle region in the original micrograph after normalization (Figure 5(b)), the particles in Figure 5(c) and (k) has more intensity contrast and are more isolated from the background than the ones in Figure 5(a) and (b), which will make it easier for clustering algorithms to identify them.

**Figure 5.**
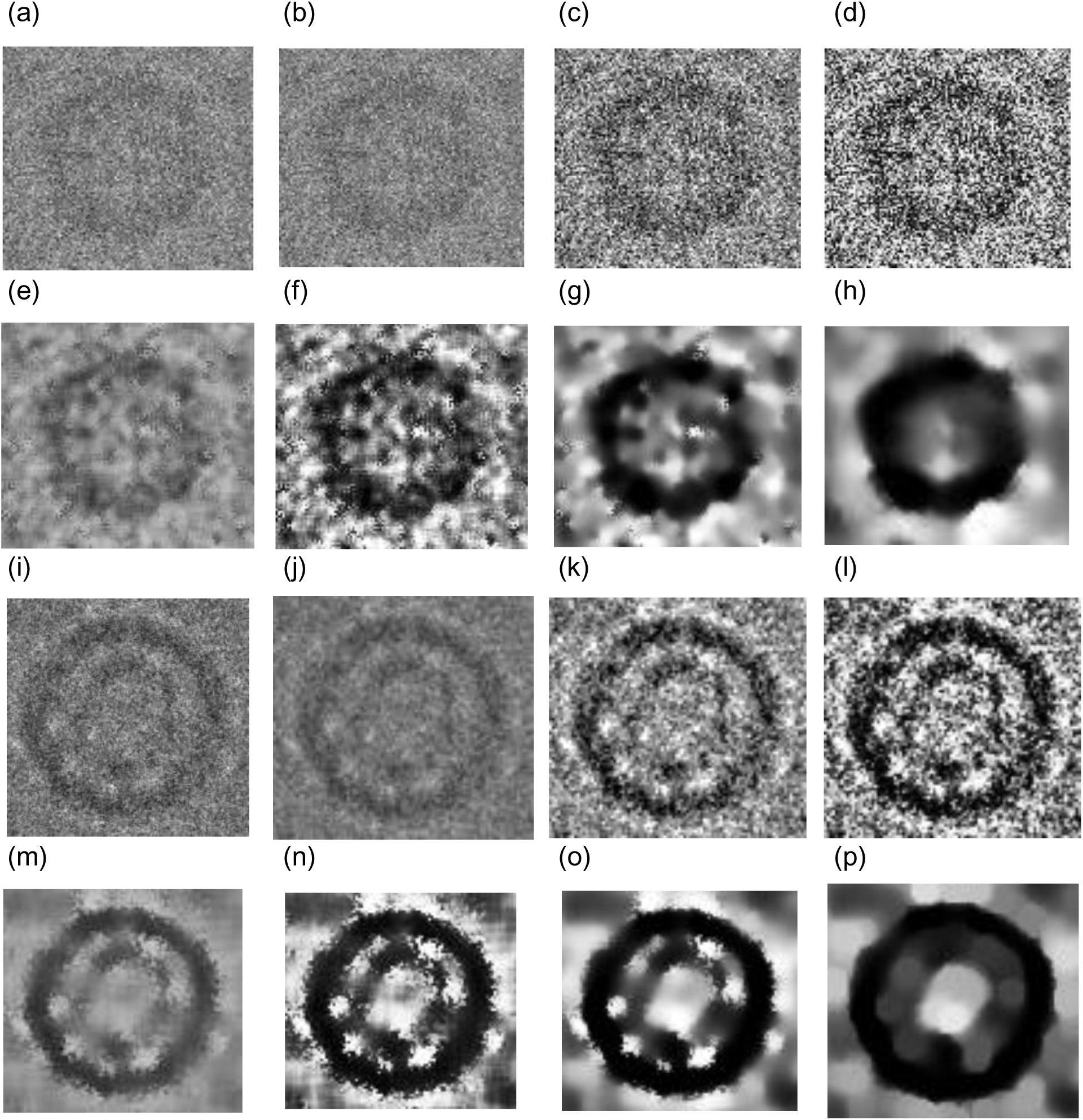
Illustration of effects of the cryo-EM image analysis on a zoom-in selected particle region using two different examples from two datasets. (a) An original zoom-in selected particle region in the micrograph image in Apoferritin dataset, (b) The normalized single particle image region, (c) The single particle region after applying the contrast enhancement correction (CEC), (d) The single particle region after applying the histogram equalization, (e) The single particle region after applying image resonation with Wiener filtering, (f) The single particle region after applying the contrast-limited adaptive histogram equalization, (g) The single particle region after applying image guided filtering, (h) The single particle region after applying morphological image operation, (i) An original zoom-in selected particle region in a micrograph image in the KLH dataset before the preprocessing steps, (j) The selected particle region in a micrograph image in the KLH dataset after normalization, (k) The selected particle region in a micrograph image in the KLH dataset after applying the contrast enhancement correction (CEC), (l) The selected particle region in a micrograph image in the KLH dataset after applying the histogram equalization, (m) The selected particle region in a micrograph image in the KLH dataset after applying image resonation with Wiener filtering, (n) The selected particle region in a micrograph image in the KLH dataset applying the contrast-limited adaptive histogram equalization, (o) The selected particle region in a micrograph image in the KLH dataset after applying image guided filtering, (p) The selected particle region in a micrograph image in the KLH dataset after applying morphological image operation.

### Step3: Global Cryo-EM Contrast Enhancement

Due to the low-dose micrograph imaging mod on a large defocuses particles area, cryo-EM images have low contrast areas where the particles are difficult to detect. Histogram equalization [29] based on a uniform distribution is used to increase and enhance the intensity value of the image pixels. It increases and improves the global image contrast by mapping the original image histogram to a uniform histogram. Figure 5(d) and (l) show an example of a selected particle region in the micrograph after global contrast enhancement-based histogram equalization. Compared with the pervious step (e.g. Figure 5(c) and (k)), the particle object regions have more contrast with the background.

### Step 4: Cryo-EM Noise Suppressing

Due to the small electron doses and low contrast between protein and solvent, cryo-EM images tend to be rather noisy [30]. Image restoration is applied to denoise single particle cryo-EM images [31]. Based on the prior knowledge of the degradation process, the image restoration recovers and improves the quality of an image by identifying the type of noise and then removing it. Since the cryo-EM images are often corrupted by typically gaussian noise, the Weiner filter is chosen to model the noise. The Wiener filter is applied to remove additive noise and invert the blurring in cryo-EM images [32]. It minimizes the overall mean square error in the process of inverse filtering and noise smoothing. The Wiener filter in the Fourier domain can be expressed as in Equation (3).

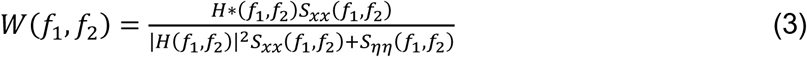

Where *S*_*xx*_ (*f*_1_,*f*_2_) + *S_*ηη*_*(*f*_1_,*f*_2_) are respectively the power spectra of the original image and the additive noise, and *H*(*f*_1_, *f*_2_) is the blurring filter. Figure 5(e) and (m) show two different zoom-in particles after applying noise suppressing based image restoration using Wiener filtering. We can notice that, in both cases, some background noise is removed, and the structure of the particle object appears more distinctly than the particle object in the previous step (Figure 5(a)-(d)).

### Step 5: Local Particles Contrast Enhancement in cryo-EM

In general, particle picking process depends basically on the quality of the particles in the cryo-EM. Since there are too many low-quality particle shapes in the cryo-EM images, the local features of the particles such as the contrast, intensity level, and edges, need to be improved and enhanced [26]. In term of locally enhance the particle edges in the cryo-EM, adaptive histogram equalization (AHE) [32] is used to improve the local contrast in images. It provides a sophisticated technique for contrast dynamic range modification based on the intensity histogram shape description. It is applied to small regions of cryo-EM images, called tiles. It enhances the contrast of each tile so that the histogram of the output region approximately matches a specified histogram. The Adaptive Histogram Equalization combines neighboring tiles using bilinear interpolation to eliminate artificially induced boundaries. It is based on a probability model to enhance the contrast condition of each small region (sub-rejoin) using Equation (4) [32]:

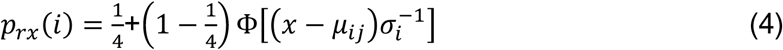

where *p_rx_*(*i*) is the image contrast-limited adaptive histogram equalization function of pixel value and Ф denotes the cumulative gaussian distribution function for each region, which has a separate location parameter estimate for each region. 1/4 is a constant for the 4-choice task [32]. Figure 5(f) and (n) show two different zoom-in particles after applying local particles contrast enhancement based on contrast-limited adaptive histogram equalization. The particle object intensity (contrast) is significantly improved and enhanced. In both examples, particles look darker and have a higher contrast than the previous particle images (Figure 5(e) and (m)).

### Step 6: Particle Edges Enhancement in cryo-EM

In order to localize each particle object in the cryo-EM image, particle edges enhancement is proposed to isolate the particle shapes in the cryo-EM image. Edge-preserving smoothing technique is used to locally smooth and enhance the particle edges in order to localize different particles in any cryo-EM. Guided image filtering [33] is employed to perform edge-preserving and smoothing using the content of a second image, called a guidance image, to influence the filtering. The guided filter generates the filtered output by considering the content of a guidance image, which can be the input image itself or a different image. It has a theoretical connection with the matting Laplacian matrix [33] and can better utilize the structures in the guidance image. Let us assume that *I* is a guidance image filter, *p* is an input cryo-EM image, and *q* is an output image. Both *I* and *p* are given beforehand and can be identical. The filtered output at a pixel *i* is expressed as a weighted average as shown in Equation (5) [33]:

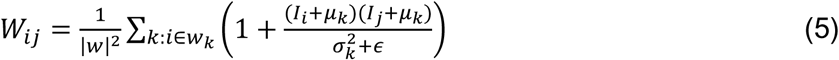

where *i* and *j* are pixel indices. The filter kernel *W*_*ij*_ is a function of the guidance cryo-EM image *I* and independent of *p* as in Equation (6) [33]:

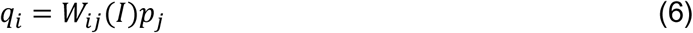

where *q*_*i*_ is the output image after the image guidance filtering and *p*_*j*_ is the input image after the image guidance filtering. A MATLAB function “*imguidedfilter*” is used to implement the guided filtering. It performs the edge-preserving smoothing of the cryo-EM image in order to reduce the noise while keeping the particle edges. Figure 5(g) and (o) show two different zoom-in particles after applying particle edges enhancement using image guided filtering. The overall contrast of the particle in the cryo-EM image is improved. Compared to the same particle in the previous step (Figure 5(f) and (n)), particle edges appear more smoothly and some dark spots around the particle object become smoother and brighter while particle object edges become darker. In addition, the particle edges are more connected and have higher contrast than the background.

### Step 7: Particle Shape Localization in cryo-EM

The last step of the pre-processing stage is the particle object localization and isolation step. In this step, we use a morphological image processing [29], which is a collection of non-linear operations related to the shape or morphology of features in an image. Logical operations are applied to make particle regions similar to each other and different from the background regions. We apply an opening dilation operation followed by erosion with the same structuring element as shown in Equation (7) [29]:

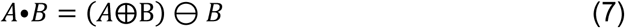

where *A* is the original cryo-EM image and *B* is the structure element. Figure 5 (h) and (p) show two different zoom-in particles after applying shape localization using morphological image operation (image closing with a structural element 5×5). The particle object is significantly improved and more isolated from the background. Also, the particle object structure is fully connected and has a higher contrast. The particle background is smother, compared to the particle background in the previous step Figure 5(g) and (o).

### Stage 2: Particle Clustering

In this stage, a binary mask is constructed using unsupervised learning clustering methods for particle isolation. Two standard clustering algorithms K-means [23] and FCM) [24] as well as a new intensity-based clustering (IBC) algorithm are applied. This clustering algorithm is based on an intensity distribution model, *P*(*i*; *d*), which relates the intensity difference value *d* to signed difference intensity values, *i*.

Let assume that {*I*_2_,*I*_2_,*I*_*n*_} be a set of images of the same modality containing the same anatomical structure of various subjects (i.e. particles in the cryo-EM images), and {*x*^(1)^, *x*^(2)^, *x*^(*L*)^ } be all (L) the pixels in an image to be grouped into several consistent “clusters” where the number of cluster is determined according to a specific intensity interval size. To determine the initial number of cluster in the ICB algorithm, for example *K* = 4, if the adjusted intensity range is [0.2, 0.8] as shown in Figure 3(b) and the interval size is 0.15, there are 4 initial cluster levels: the intensity level [0.2-0.35] will be assigned to Cluster 1, [0.35-0.5] to Cluster 2, [0.5-0.65] to Cluster 3, and [0.65-0.8] to Cluster 4. Here, *x*^(*i*)^ is a real intensity value in a specific range, 1 <= *i* < = *L*. Let {*θ*_1_, *θ*_2_,… *θ*_*K*_} be the set of the average intensity values of *K* clusters. Let *U*_*j*_ be the index of the cluster whose center (*θ*_*j*_) is closest to *x*^(*i*)^. The cluster assignment of all pixels 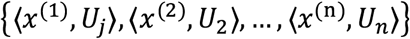 is updated iteratively according to the average intensity (*θ*_*j*_) of clusters. The centers (*θ_j_*) of K clusters are initialized as evenly distributed intervals in the intensity range at the equal step size according to Equation (8):

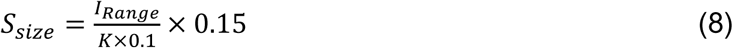

where the *I*_*Range*_ is the difference between the maximum and the minimum intensity level in each image and 0.15 is the selected interval step size value. The procedure of the clustering algorithm for cryo-EM image clustering is shown below.

#### Algorithm 1 Intensity Based Clustering (IBC)

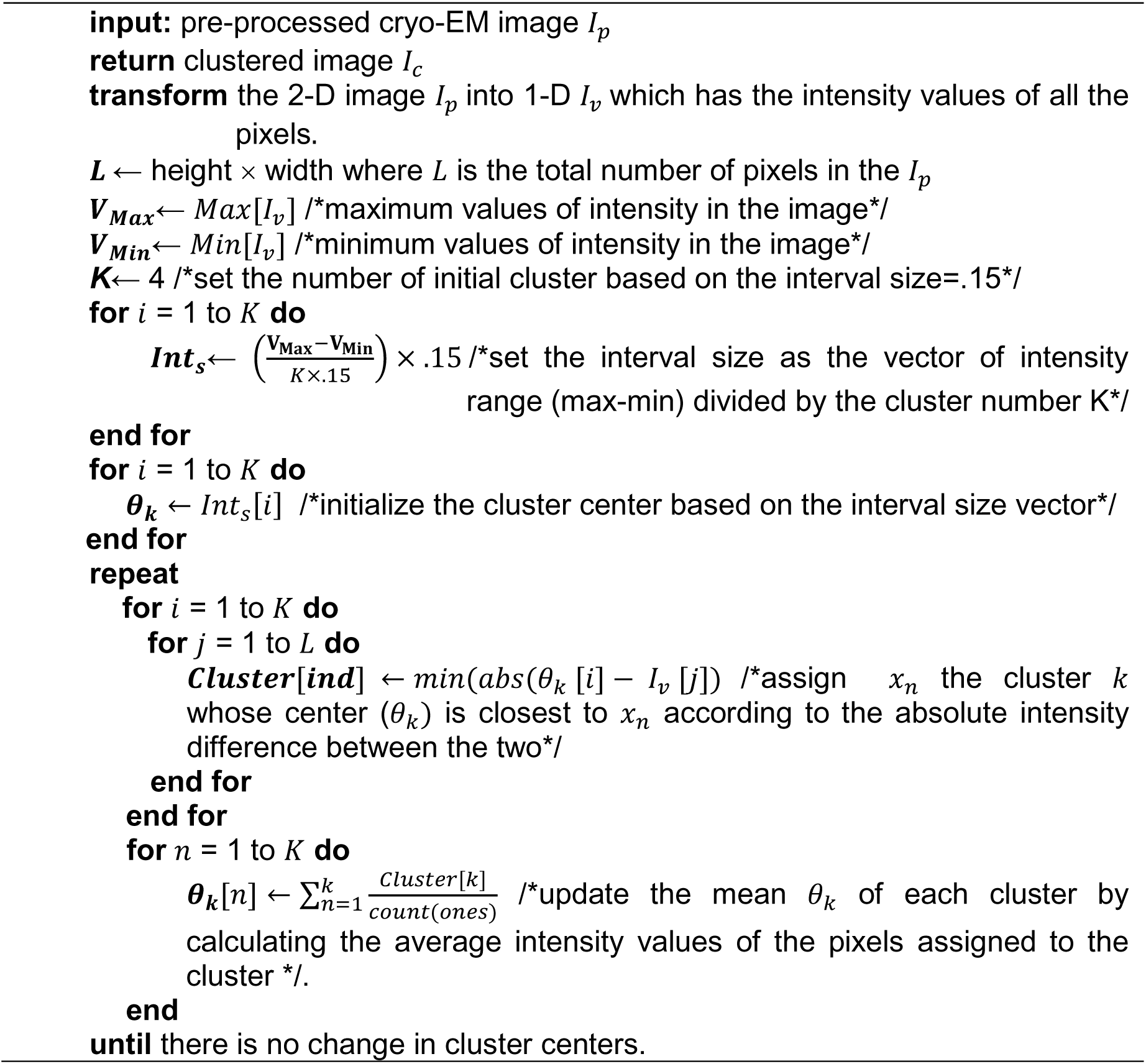

Figure 6(a) and (b) show an example of different cryo-EM clustering results by using the intensity-based clustering method (ICB) with two cryo-EM datasets (Apoferritin [34] and KLH datasets [35]).

**Figure 6.**
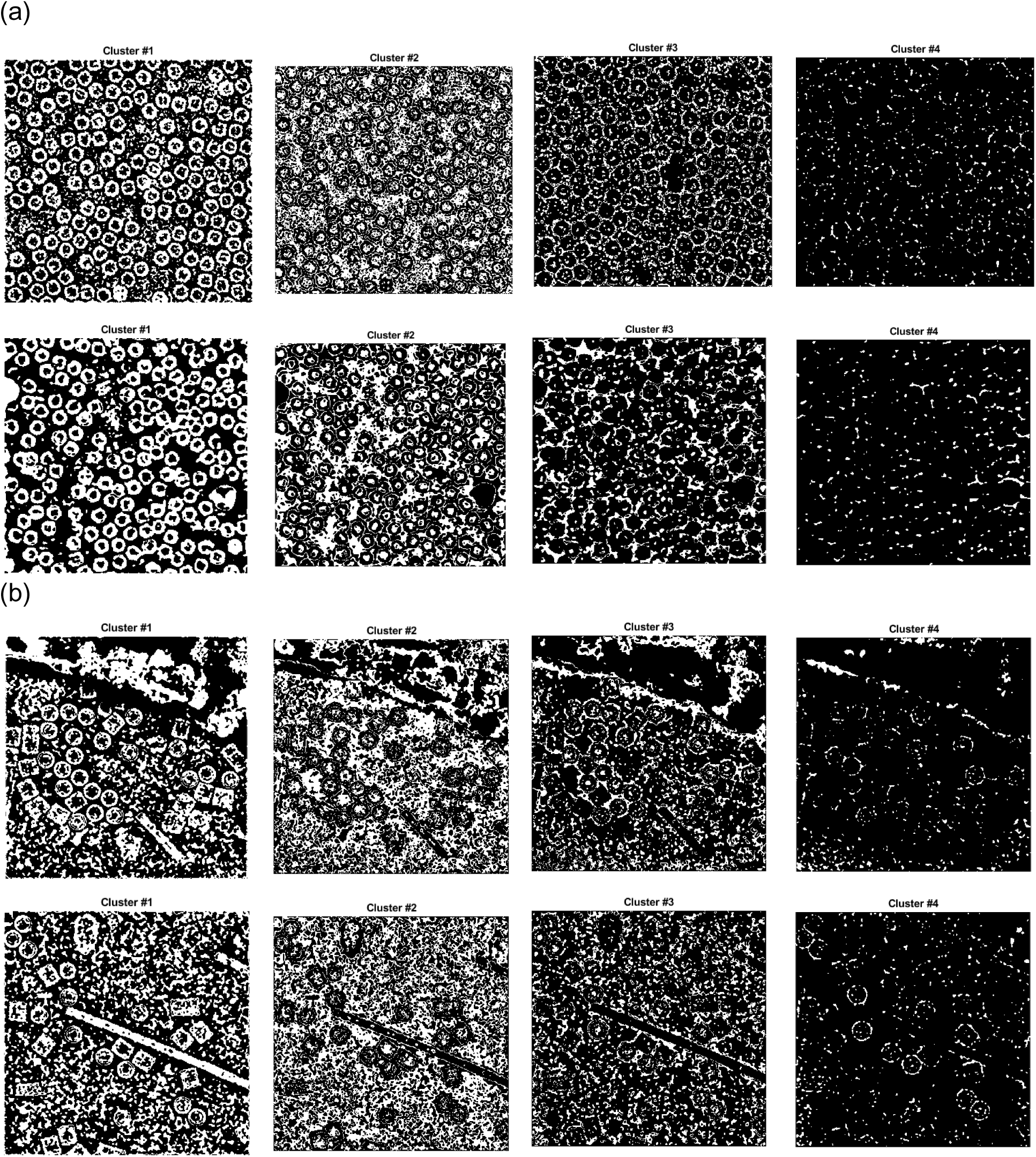
Different cryo-EM image clustering results using an Intensity-Based Clustering Algorithm (ICB). (a) Two sets of cryo-EM image clustering results (Cluster #1, Cluster #2, Cluster #3 and Cluster #4) on the Apoferritin dataset. Most real particles were always assigned to Cluster 1, (b) Two sets of cryo-EM image clustering results (Cluster #1, Cluster #2, Cluster #3 and Cluster #4) on the KLH dataset. Most real particles were always assigned to Cluster 1.

It is noticed that the particles are most stably grouped in Cluster 1. Generally, the particles of the different images of the same protein can be best identified in the same specific cluster by the ICB method according to our experiments. However, the particles are not most stably grouped in the same cluster by k-means and FCM algorithms due to their random initialization of cluster centers. Figure 7 and 8 show the clustering results of the same cryo-EM images using k-means and FCM respectively.

**Figure 7.**
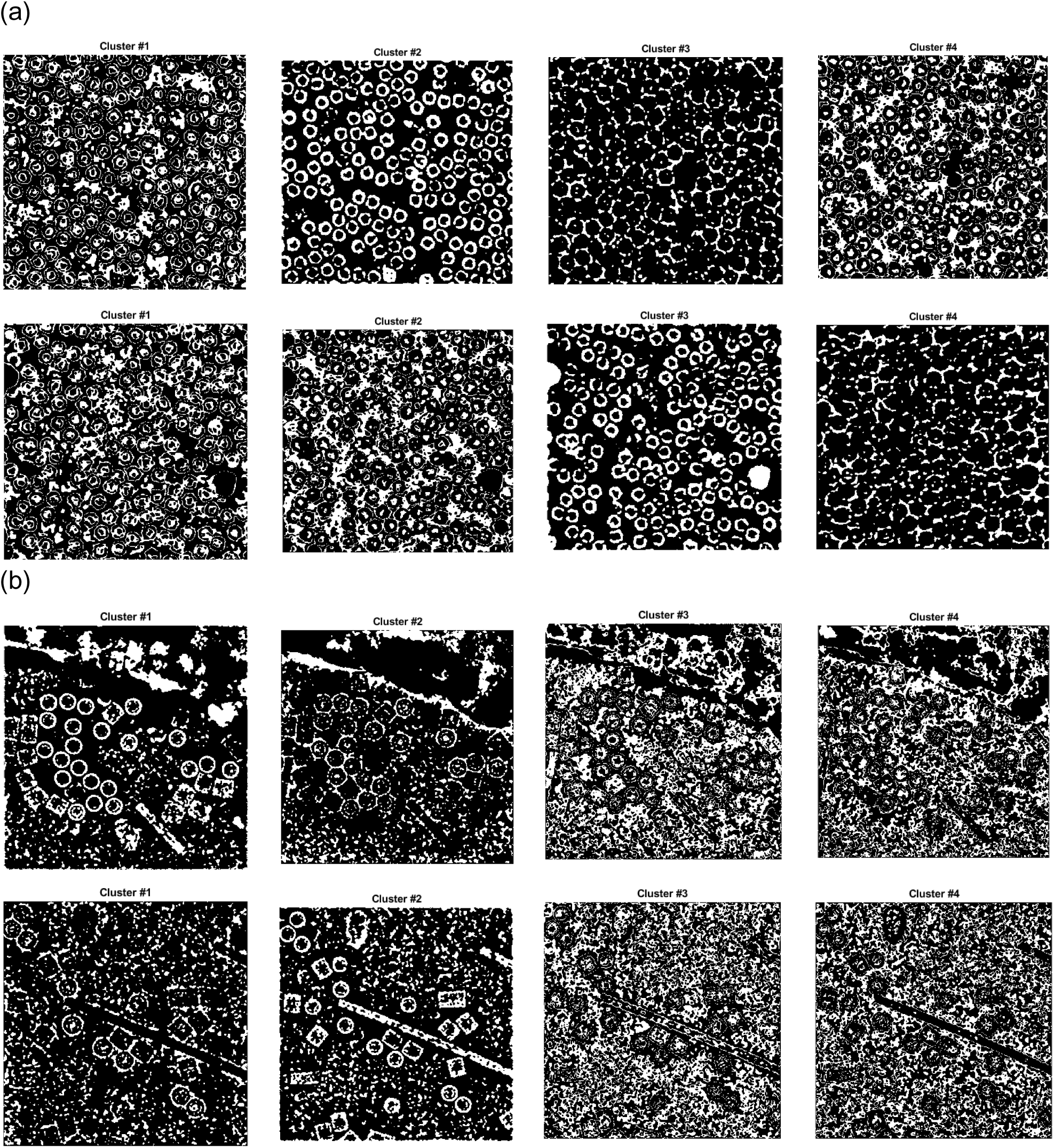
Different cryo-EM image clustering results using the k-means clustering algorithm. (a) The two sets of cryo-EM images clusters results (Cluster #1, Cluster #2, Cluster #3 and Cluster #4) on the Apoferritin dataset. Most real particles were assigned to Cluster 2 and Cluster 3, respectively, (b) The two sets of cryo-EM image clustering results (Cluster #1, Cluster #2, Cluster #3 and Cluster #4) on the KLH dataset. Most real particles were assigned Cluster 1 and Cluster 2, respectively.

It is noticed that the particles are located in different clusters. For instance, the particles clustering for two cryo-EM images in the first dataset (Apoferritin) using k-means is shown in Figure 7(a). The particles are grouped in two different clusters (Cluster 2 and 3, respectively). Figure 7(b) shows the same issue for the k-means on the second dataset (KLH). The same problem happens to FCM (Figure 8).

**Figure 8.**
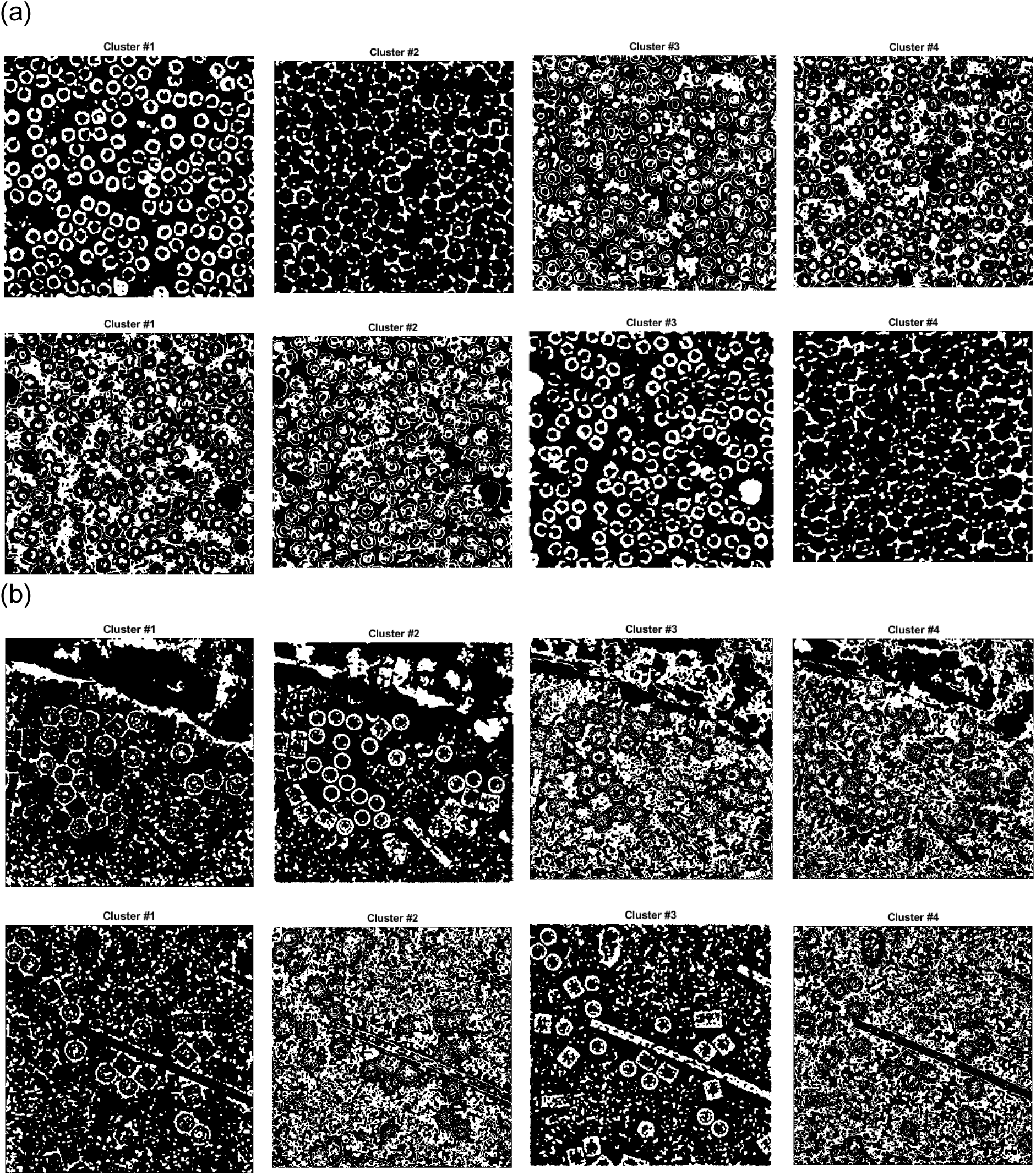
Different cryo-EM image clustering results using the FCM clustering algorithm. (a) Two sets of cryo-EM images clustering results (Cluster #1, Cluster #2, Cluster #3 and Cluster #4) on Apoferritin dataset. Most real particles were assigned to Cluster 1 and Cluster 3, respectively. (b) Two sets of cryo-EM image clustering results (Cluster #1, Cluster #2, Cluster #3 and Cluster #4) on the KLH dataset. Most real particles were assigned to Cluster 2 and Cluster 3, respectively.

### Stage 3: Particle Picking

The last stage of the AutoCryoPicker framework has two main steps. The first step is binary mask image cleaning and the second step is particle object detection and picking. In the first step, some post-processing operations (e.g. binary image region and hole filling, morphological image operation using image opening, and small object removal from the binary image) are performed to clean the binary mask produced in the clustering stage. In the second step, a modified Circular Hough Transform algorithm (CHT) [36] is applied to detect particles in the cleaned binary mask.

### Step 1: Cryo-EM Cluster Image Cleaning and Non-Circular Object Removal

A binary mask of each cryo-EM cluster image is cleaned based on removal of the small and non-circular objects via size filtering and roundness filtering. The image cleaning and small object removal algorithm is shown below.

#### Algorithm 2 Image Cleaning and Non-Circular Object Removal

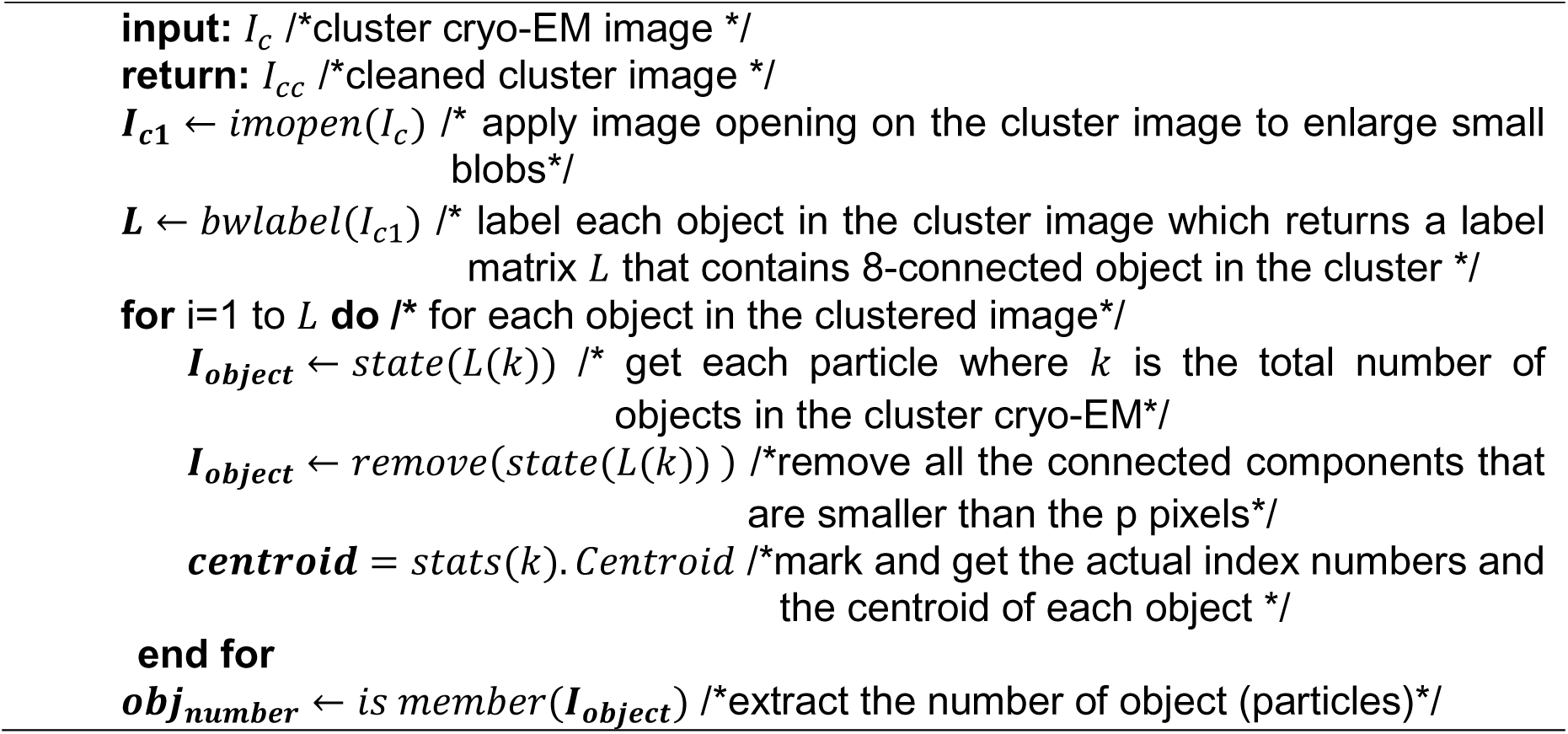
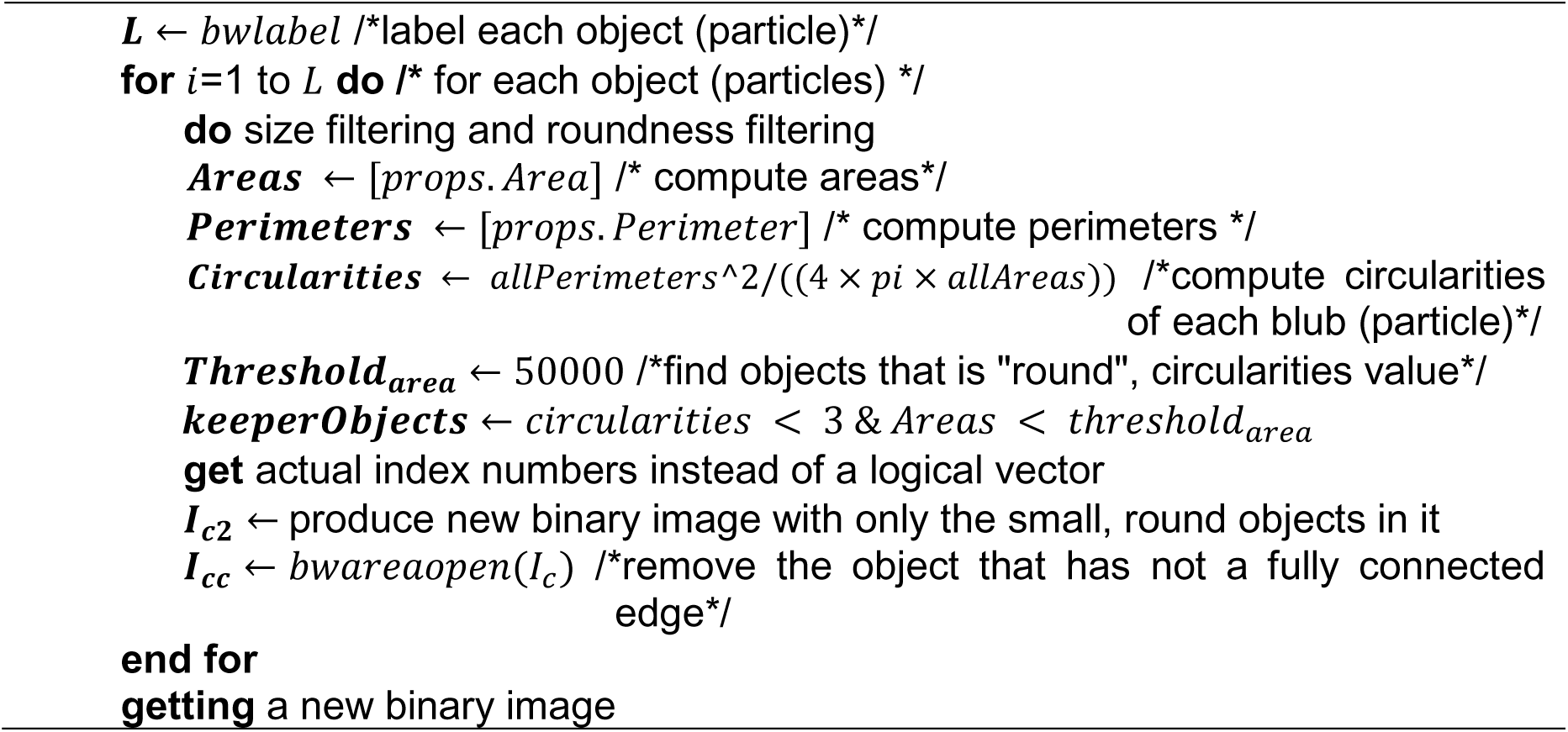

Figure 9 shows the cryo-EM image cluster cleaning results (particles clustering) before and after image cleaning step.

**Figure 9.**
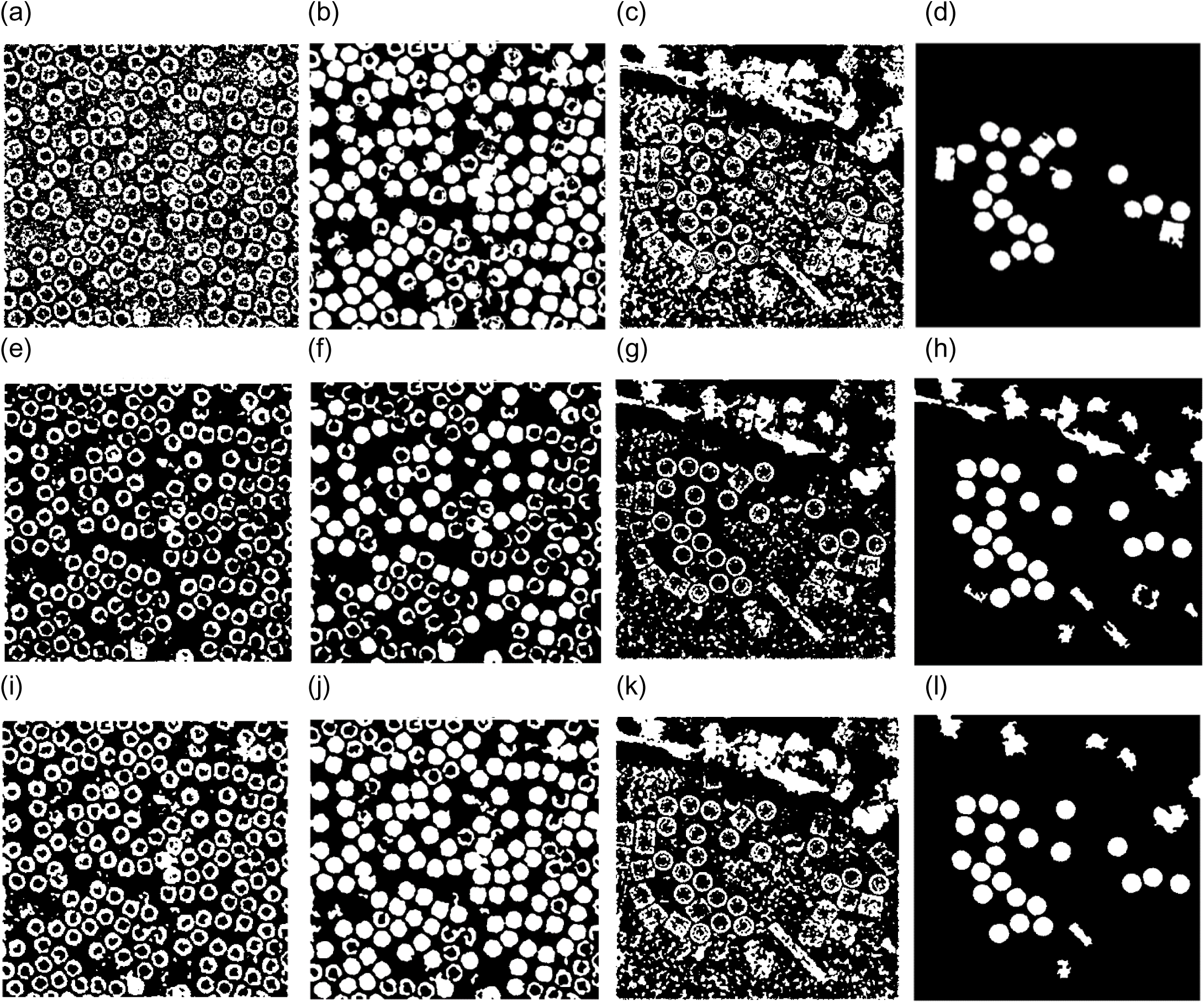
Cryo-EM Particle Clustering Results after Binary Image Cleaning and Non-Circular Object Removal. (a) The particle clustering image before binary image cleaning and non-circular object removal on the results of ICB clustering of a cryo-EM image from Apoferritin dataset, (b) The particle clustering image after binary image cleaning and non-circular object removal on the results of ICB clustering of a cryo-EM image from Apoferritin dataset, (c) The particle clustering image before binary image cleaning and non-circular object removal on the results of ICB clustering of a cryo-EM image from KLH dataset, (d) The particle clustering image after binary image cleaning and non-circular object removal on the results of ICB clustering of a cryo-EM image from KLH dataset, (e) The particle clustering image before binary image cleaning and non-circular object removal on the results of k-means clustering of a cryo-EM image from Apoferritin dataset, (f) The particle clustering image after binary image cleaning and non-circular object removal on the results of k-means clustering of a cryo-EM image from Apoferritin dataset, (g) The particle clustering image before binary image cleaning and non-circular object removal on the results of k-means clustering of a cryo-EM image from KLH dataset, (h) The particle clustering image after binary image cleaning and non-circular object removal on the results of k-means clustering of a cryo-EM image from KLH dataset, (i) The particles clustering image before binary image cleaning and non-circular object removal on the results of FCM clustering of a cryo-EM image from Apoferritin dataset, (j) The particle clustering image after binary image cleaning and non-circular object removal on the results of FCM clustering of a cryo-EM image from Apoferritin dataset, (k) The particle clustering image before binary image cleaning and non-circular object removal on the results of FCM clustering of a cryo-EM image from KLH dataset, (l) The particle clustering image after binary image cleaning and non-circular object removal on the results of FCM clustering of a cryo-EM image from KLH dataset.

Figure 9(b), (f), and (j) show the particles clustering and cleaning results for the cryo-EM images form the Apoferritin dataset using ICB, k-means, and FCM respectively. Figure 9(d), (h), and (I) show the particles clustering and cleaning results for the cryo-EM images form the KLH dataset using ICB, k-means, and FCM respectively. It noticed that the proposed algorithm (ICB) produces significantly cleaner clustering images than the other two standard clustering algorithms.

### Step 2: Top View (Circular) Particle Detection and Picking in Cryo-EM

Since the regular shape of the protein particle in the test cryo-EM dataset is a common shape – circle (top view), a Circular Hough Transform (CHT) is used to detect particles in cluster images. For another common particle shape in cryo-EM images - square, a square shape detector would be needed.

To detect each circular particle in every cryo-EM image, we apply a modified Circular Hough Transform algorithm (CHT) as follows.

#### Algorithm 3 Circular Hough Transformation (CHT)

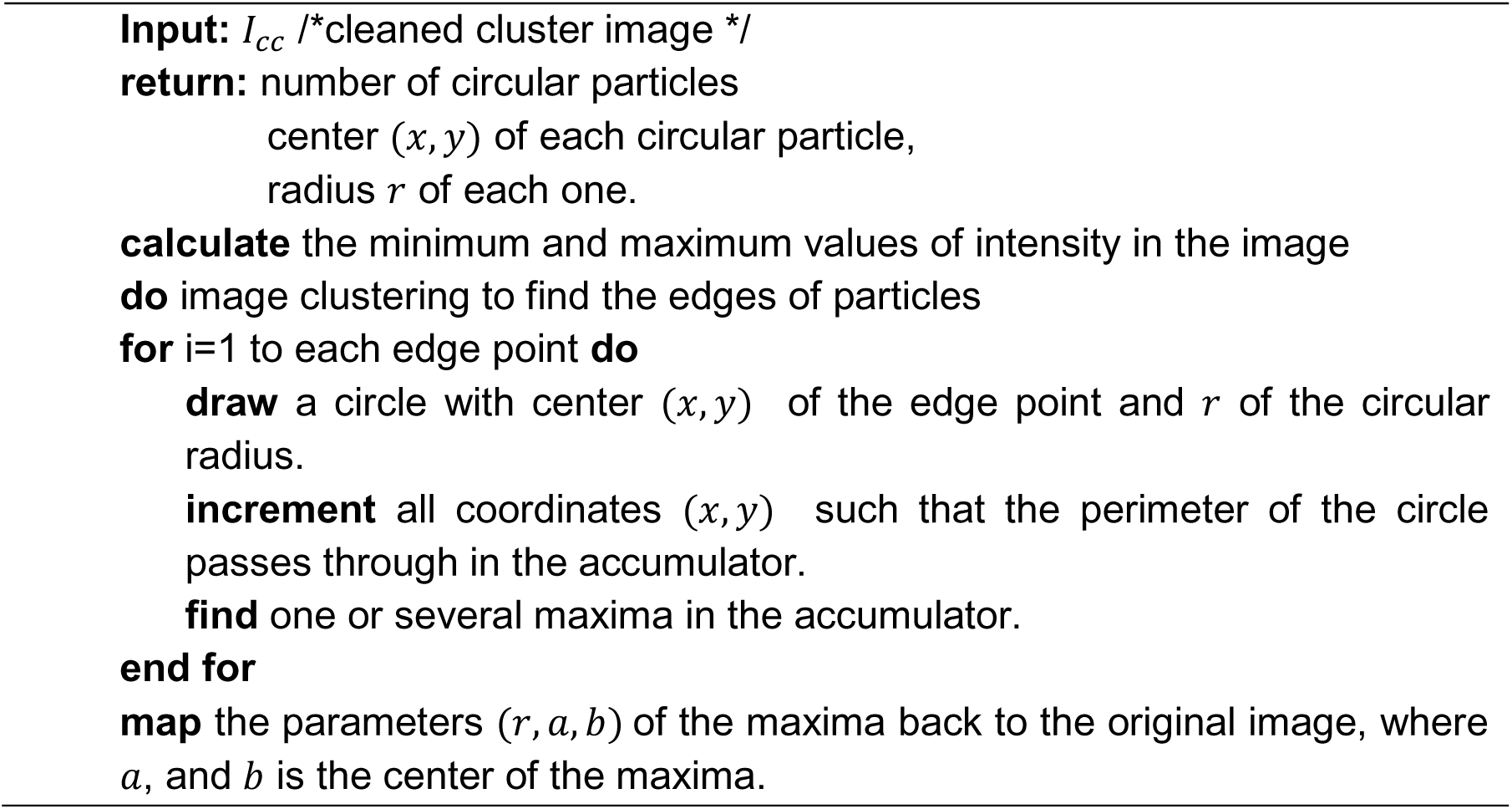

The detection algorithm returns the center and radius of each particle as is shown in Figure 10(c), (f), (i), (m), (p), and (s) based on the clustering results of the different clustering algorithms (ICB, k-means, and FCM) respectively for Apoferritin and KLH datasets. For instance, Figure 10(c) shows the center and radius of each particle illustrated by a ‘+’ sign and a blue circle. A bounding box is drawn around each particle object in the cryo-EM image (Figure 10(d)).

**Figure 10.**
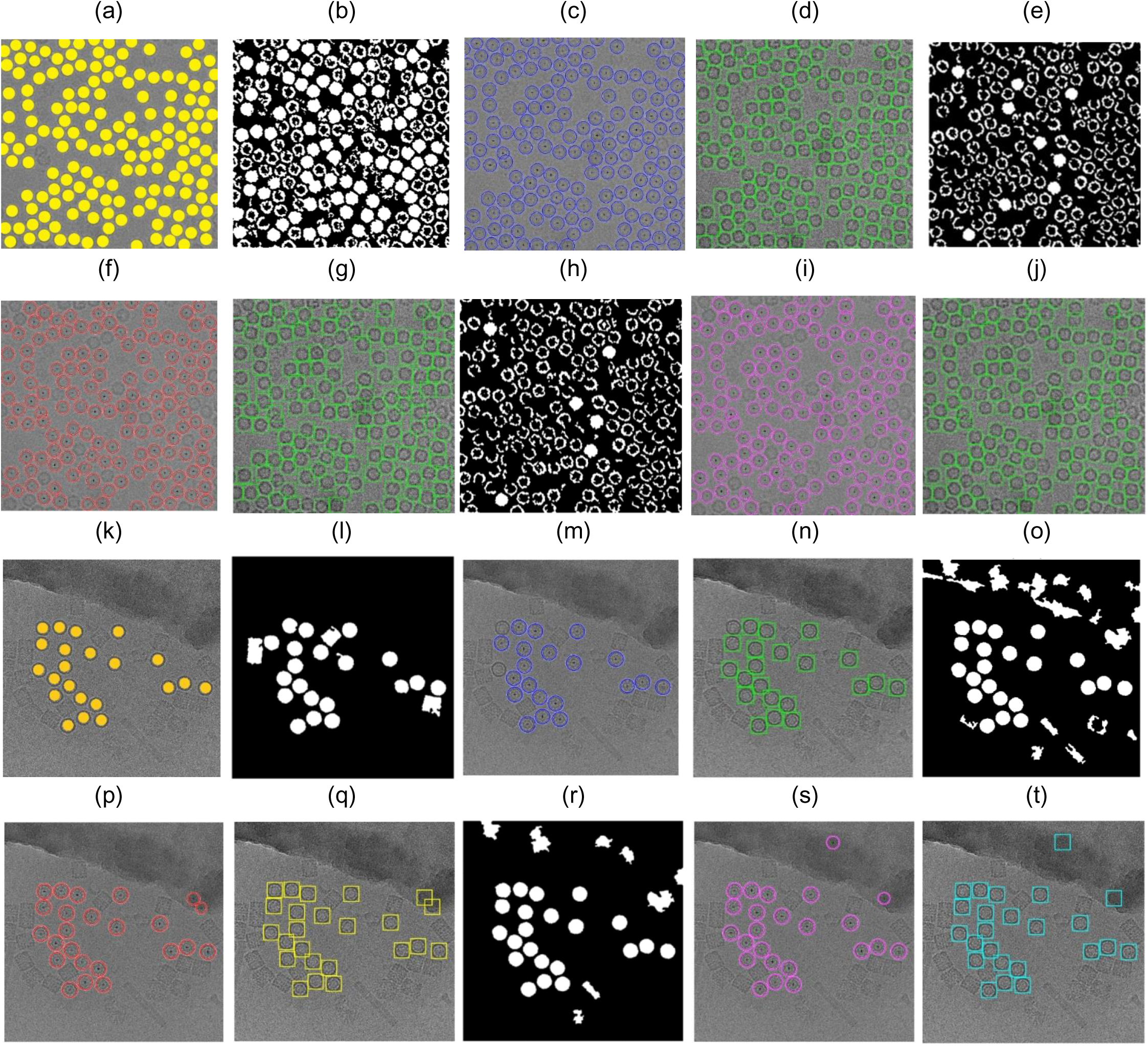
Top View (Circular) Particles Detection and Picking Results using Modified Circular Hough Transform (CHT). (a) The Ground truth (particles manually labelled) for the cryo-EM image from the Apoferritin dataset, (b) ICB clustering results after the binary image cleaning and non-circular objects removal (Apoferritin dataset), (c) The center of each particle illustrated by the ‘+’ sign and the radius of each particle by the blue circle around each particle (ICB and Apoferritin dataset), (d) The bounding box for each particle object in the original cryo-EM image (ICB and Apoferritin dataset), (e) K-means clustering results after the binary image cleaning and non-circular objects removal (Apoferritin dataset), (f) The center of each particle illustrated by using the ‘+’ sign and the radius of each particle by the blue circle around each particle (k-means results on Apoferritin dataset), (g) The bounding box for each particle (k-means results and Apoferritin dataset), (h) FCM clustering results after the binary image cleaning and non-circular objects removal (Apoferritin dataset), (i) The center of each particle illustrated by the ‘+’ sign and the radius of each particle by the blue circle around each particle (FCM and Apoferritin dataset), (j) The bounding box for each particle in the original cryo-EM image (FCM results and Apoferritin dataset), (k) The ground truth (particles manually labeled) for the cryo-EM image from the KLH dataset, (l) ICB clustering results after the binary image cleaning and non-circular objects removal (KLH dataset), (m) The center of each particle illustrated by the ‘+’ sign and the radius of each particle by the blue circle (ICB and KLH dataset), (n) The bounding box for each particle in the original cryo-EM image (ICB and KLH dataset), (o) K-means clustering results after the binary image cleaning and non-circular objects removal (KLH dataset), (p) Shows the center of each particle illustrated by the ‘+’ sign and the radius of each particle by the blue circle (k-means and KLH dataset), (q) The bounding box for each particle in the original cryo-EM image (k-means and KLH dataset), (r) FCM clustering results after the binary image cleaning and non-circular objects removal (KLH dataset), (s) The center of each particle illustrated by the ‘+’ sign and the radius of each particle by the blue circle (FCM and KLH dataset), (t) The bounding box for each particle in the original cryo-EM image (FCM and KLH dataset).

Figure 10(c) and (d) show the results of the particle object detection and picking based on the ICB clustering and the Circular Hough Transform (CHT) on the first dataset (Apoferritin). Figure 10(m) and (n) show the same results on the second dataset (KLH dataset). Figure 10(f) and (g) show the results of the particle object detection and picking based on k-means clustering and the Circular Hough Transform (CHT) on the first dataset. Figure 10(p) and (q) show the same results on the second dataset. Finally, Figure 10(h) and (j) show the results of the particle object detection and picking based on the FCM clustering and the Circular Hough Transform (CHT) on the first dataset. Figure 10(s) and (t) show the same results on the second dataset.

### Step 3: Circular and Non-Square Object Removal for the cryo-EM Cleaned Image

Another common particle shape in the cryo-EM images is a square (side view). In this case, we add another step called circular and non-square object removal form the ICB clustering image after the cleaning and small object removal step in case of keeping the side view particle shapes (square) as follows.

#### Algorithm 4 Circular and Non-Square Object (Particles) Removal

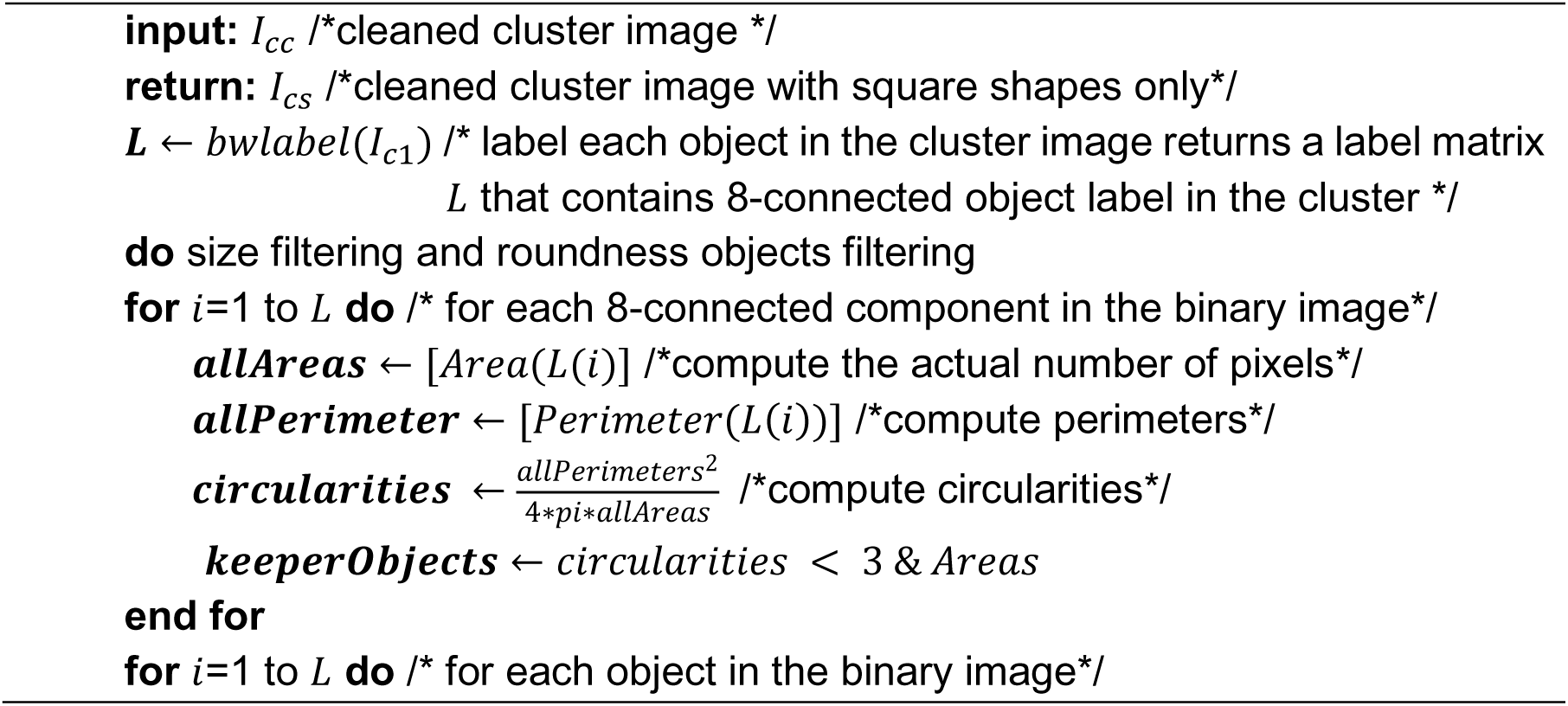
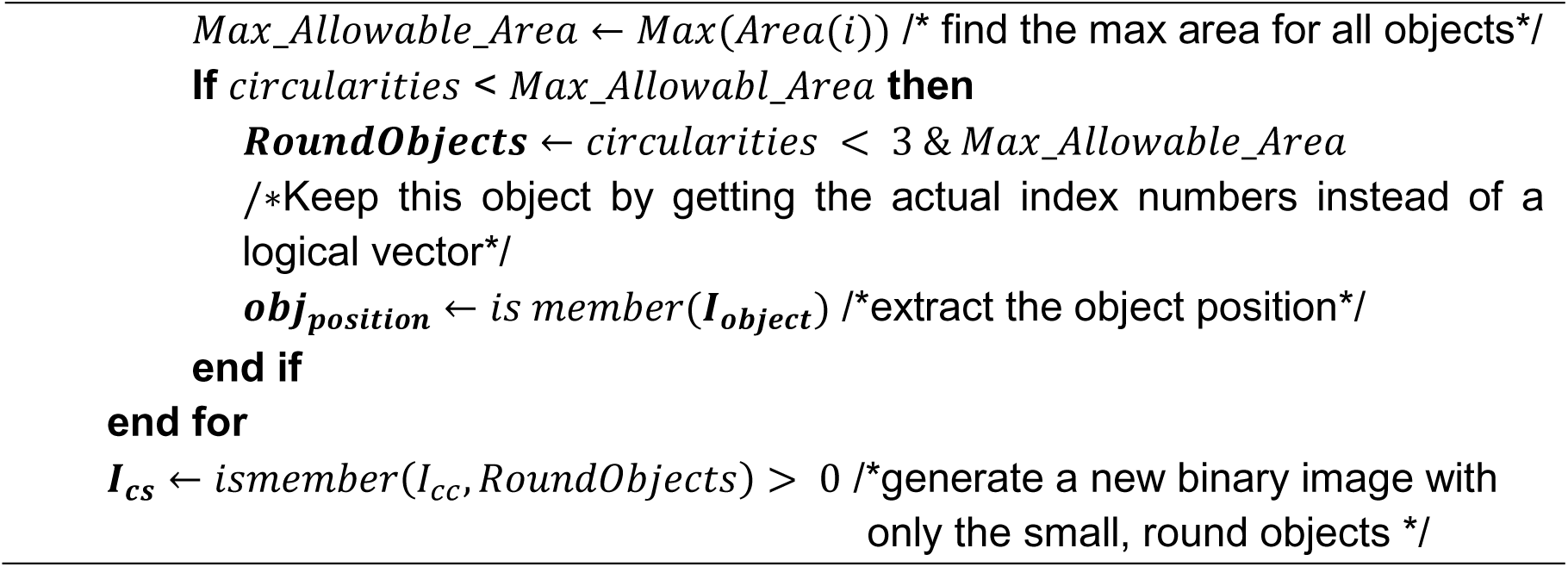

Figure 11 shows an example of the cryo-EM clustered images after the circular and non-square object removal.

**Figure 11.**
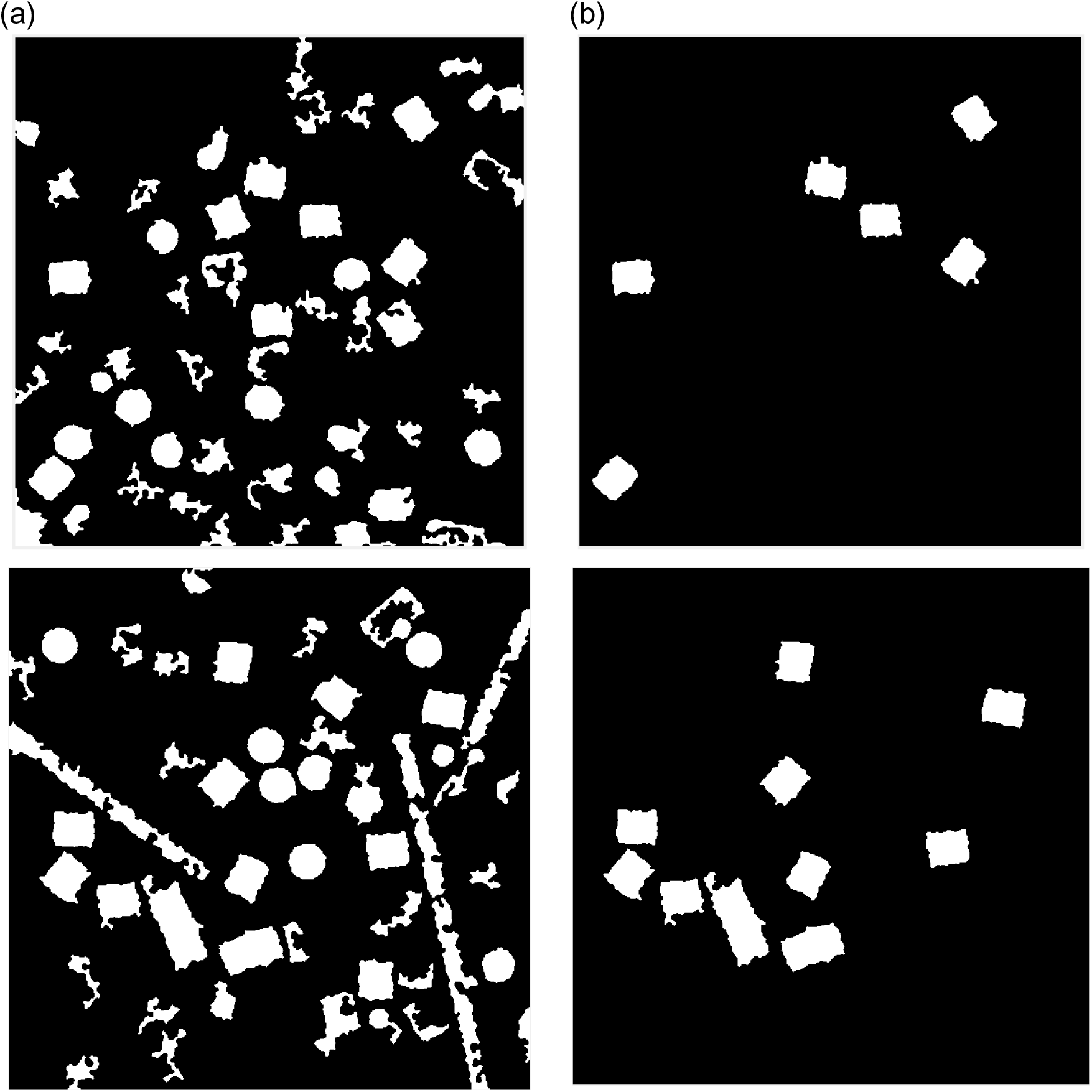
Cryo-EM clean clustered images after the circular and non-square object removal. (a) The cryo-EM clustered images after image cleaning and small objects removal, (b) The same cryo-EM clustered images after the circular and non-square object removal.

For instance, Figure 11(a) shows cryo-EM clustered images after image cleaning and small objects removal although, Figure 11(b) shows the same cryo-EM clustered images after the circular and non-square object removal. After this step, the cleaned image has only the square particle shapes (side view) in the whole cryo-EM images. We can notice that not all the particles (side view) are cleaned after the second post-processing step, but some of them are according to the similarity between the *Max_Allowable_Area* value and the circularities of each square particle object. If the circularities values between each particle shapes (side view-square and top view-circle) are very close, they are eliminated from the cleaned image.

### Step 4: Side View (Square) Particle Detection and Picking in Cryo-EM

After the circular and other object such as ice artifacts being removed, the cryo-EM cleaned mask becomes significantly clear for detecting and selecting each square particle. We apply the particle detection and picking algorithm described below.

#### Algorithm 5 Square (Side View) Particle Detection and Picking

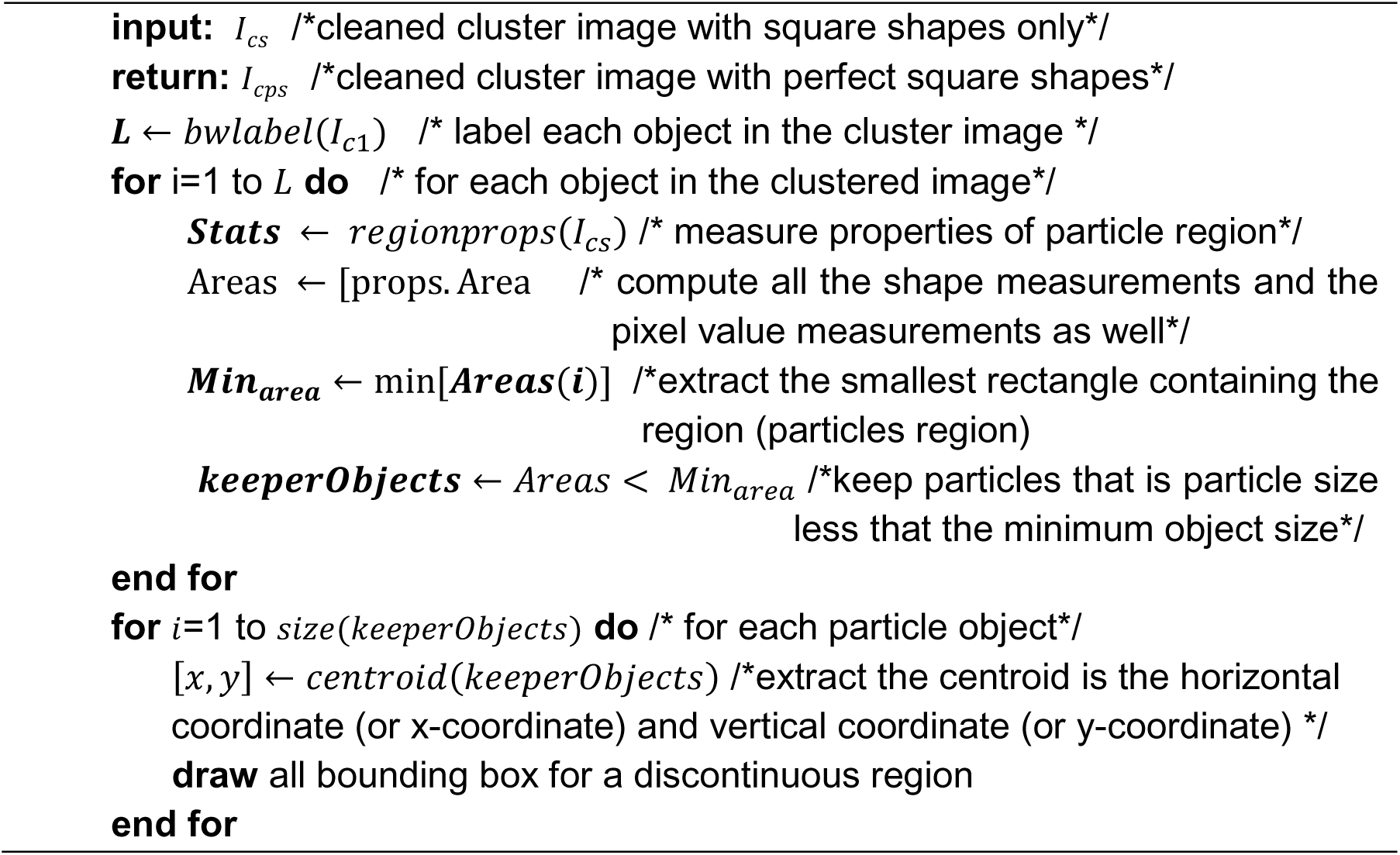

The results of the second particle shapes detection is shown in Figure 12. We can notice that some additional objects are attached to the original square particles in additional to some overlapped particles, which are also selected as shown in the final detected results in Figure 12(b).

**Figure 12.**
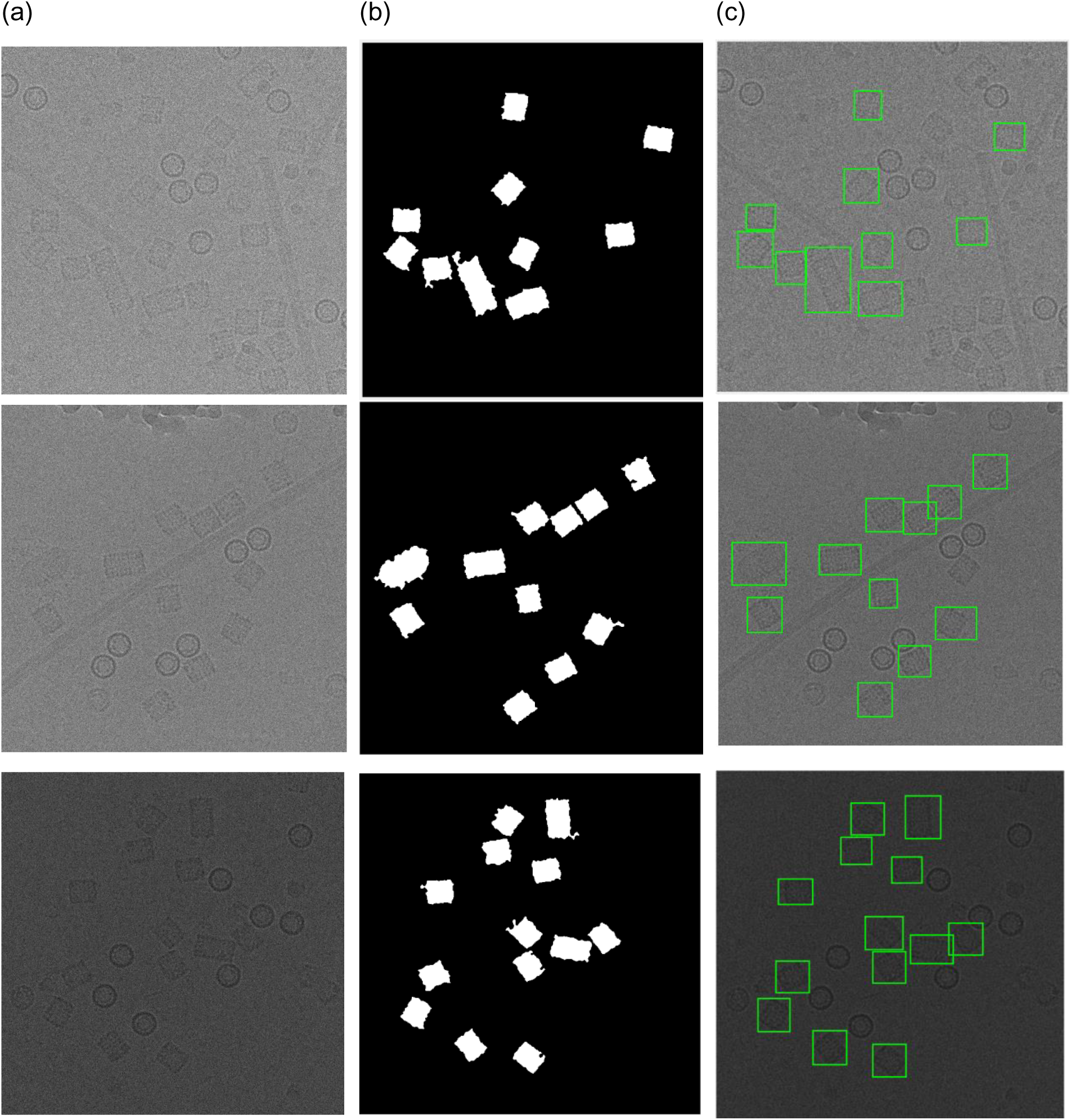
Side view (square) particles detection and picking results. (a) The original cryo-EM image (KLH dataset), (b) The result after circular and non-square object removal based on the ICB clustering algorithm, (c) Side view (square) particle detection results.

To overcome this problem, we design another postprocessing algorithm that has three main steps. The first step is applied to remove the small attached objects by blurred (smooth) each particle. Each particle convolves with a kernel (averaging filter) by using averaging filter kernel 50×50. The main particles smoothing (averaging) can be defined in Equation (9) [29]:

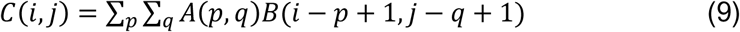

where *C*(*i*, *j*) is defined as each particle sub image, *B* is the smoothing (averaging) kernel, *p* and *q* are the particle sub image dimensions. In the second step, after the small attached object are removed for each particle, we use the Feret diameter measures approach [37] to measure and correct the particle object dimensions. New perfect particle shapes are generated based on the maximum and the minimum Feret dimeter. The maximum and minimum dimensions (width) of the particle object are used to identify the antipodal vertex pairs from the convex hull vertices set. Based on the new boundary box dimension, new perfect shapes are generated and inserted above each particle in the clean clustered cryo-EM image. The last step eliminates the outliers object (overlapped particles) by defining the average particles size and eliminating the outliers that have particle size larger than the average size. Then, the new boundary box is drawn based on the dimension of new particle object shapes.

#### Algorithm 6 Perfect Square Particles Shape Detection

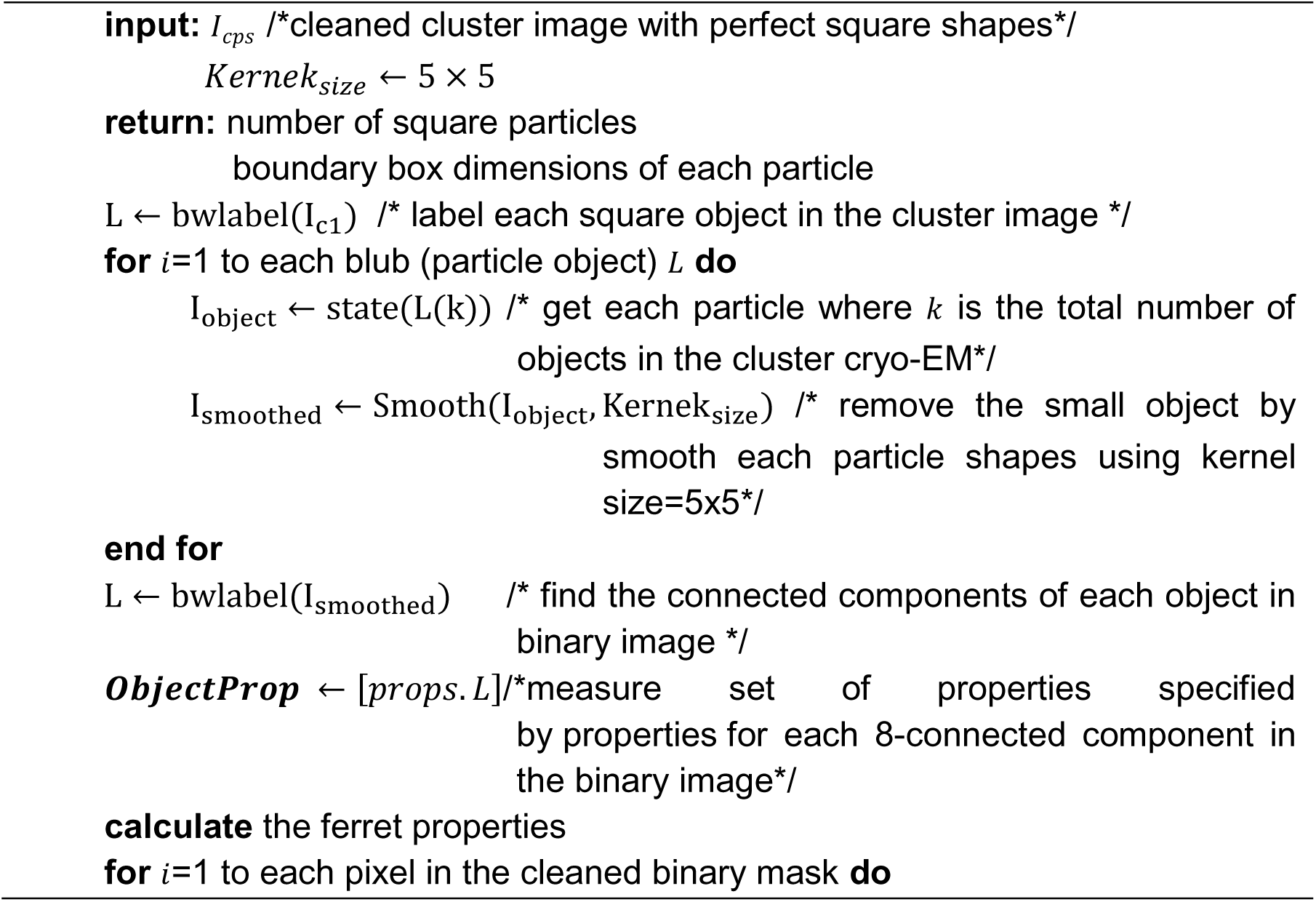
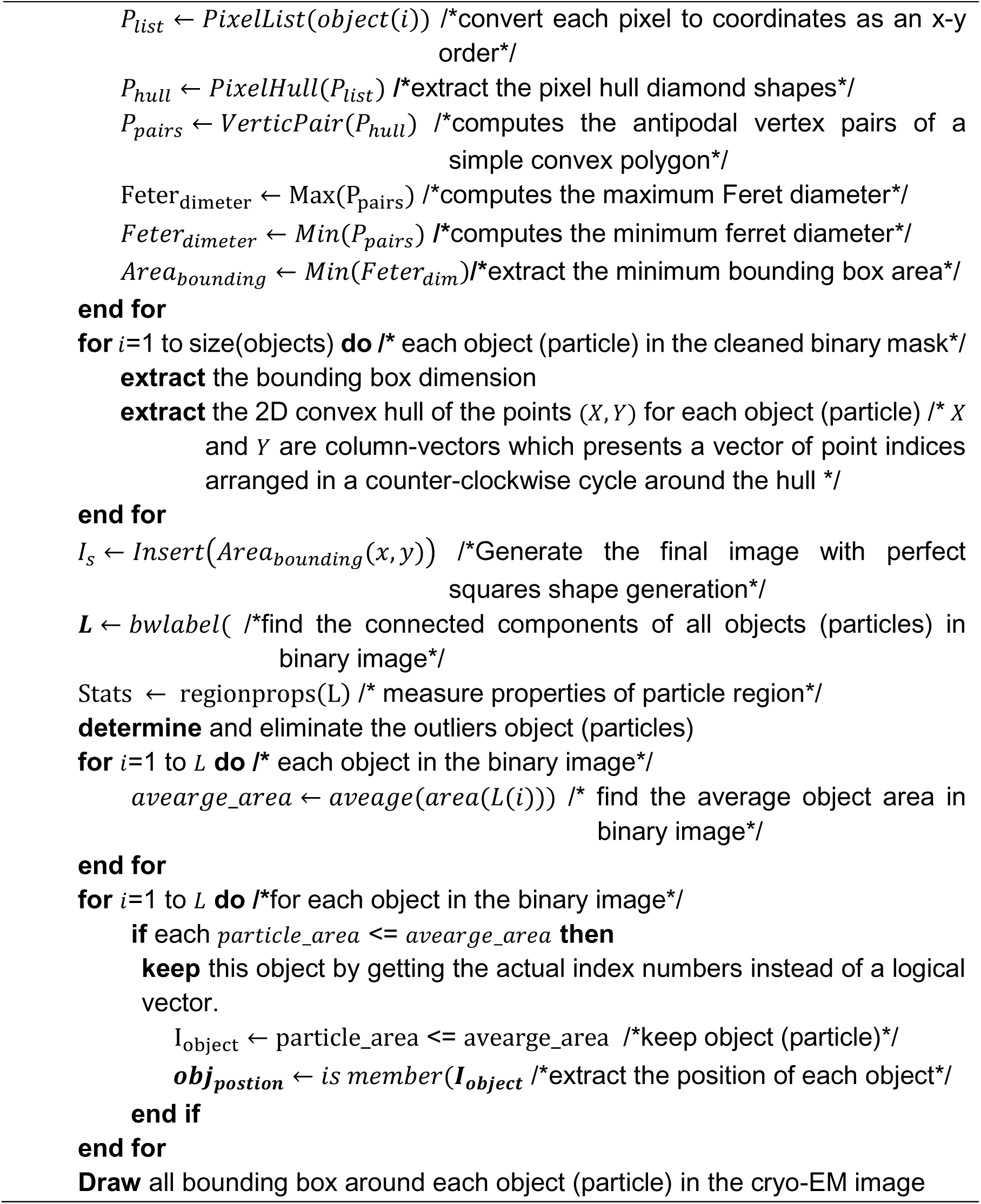

Figure 13 shows an example of the perfect square particle shapes detection using Feret object diameter. Figure 13(a) shows the square particle shapes in the image after the shapes are smoothed and blurred. Figure 13(b) shows the new boundary box of each particle based on the Feret diameter measures. Figure 13(c) shows the perfect square particle shapes based on the Feret object diameter measurement.

**Figure 13.**
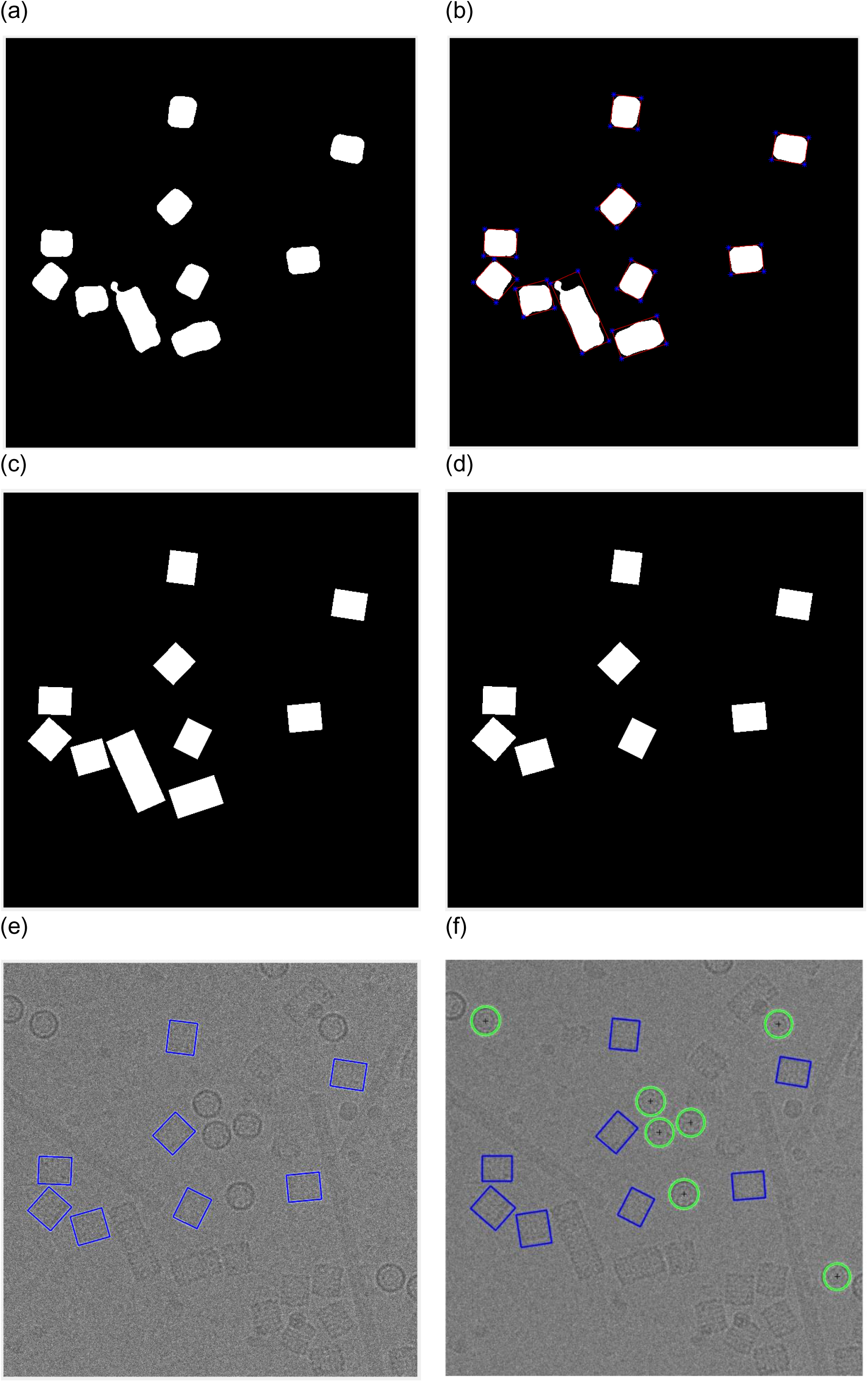
Perfect square (side view) particle shape detection using the Feret object diameter using (KLH dataset). (a) Square particle image after shapes smoothing and blurring, (b) Boundary boxes (each particle) based on Feret object diameter measurement, (c) Perfect square particle shapes that are generated based on the new boundary box dimension using Feret object diameter measurement, (d) Square particle image after the outlier objects are eliminated, (e) Square particle detection results (side view) based on the new Feret boundary box dimension, (f) The final results of two different particle shape detection and picking (top and side view) based on ICB clustering and modified CHT; and perfect square (side view) particle shapes detection using Feret object diameter.

Figure 13(d) shows the square particles image after eliminating the outlier objects (overlapped particles). Figure 13(e) shows the square particle detection results (side view) based on the new Feret boundary box. Finally, Figure 13(f) shows the final results of different particle shape detection and picking (top and side view) based ICB clustering, modified CHT, and perfect square (side view) particle shapes detection using Feret object diameter.

It is noticed that there is almost no true positive (top view particles-circle) missing. In contrast, there are some true positive example of square particles (side view) missing. Figure 14 shows some extra cases of the particle detection and picking results for both cases (top and side view) using three different algorithms (ICB, k-means, and FCM).

**Figure 14.**
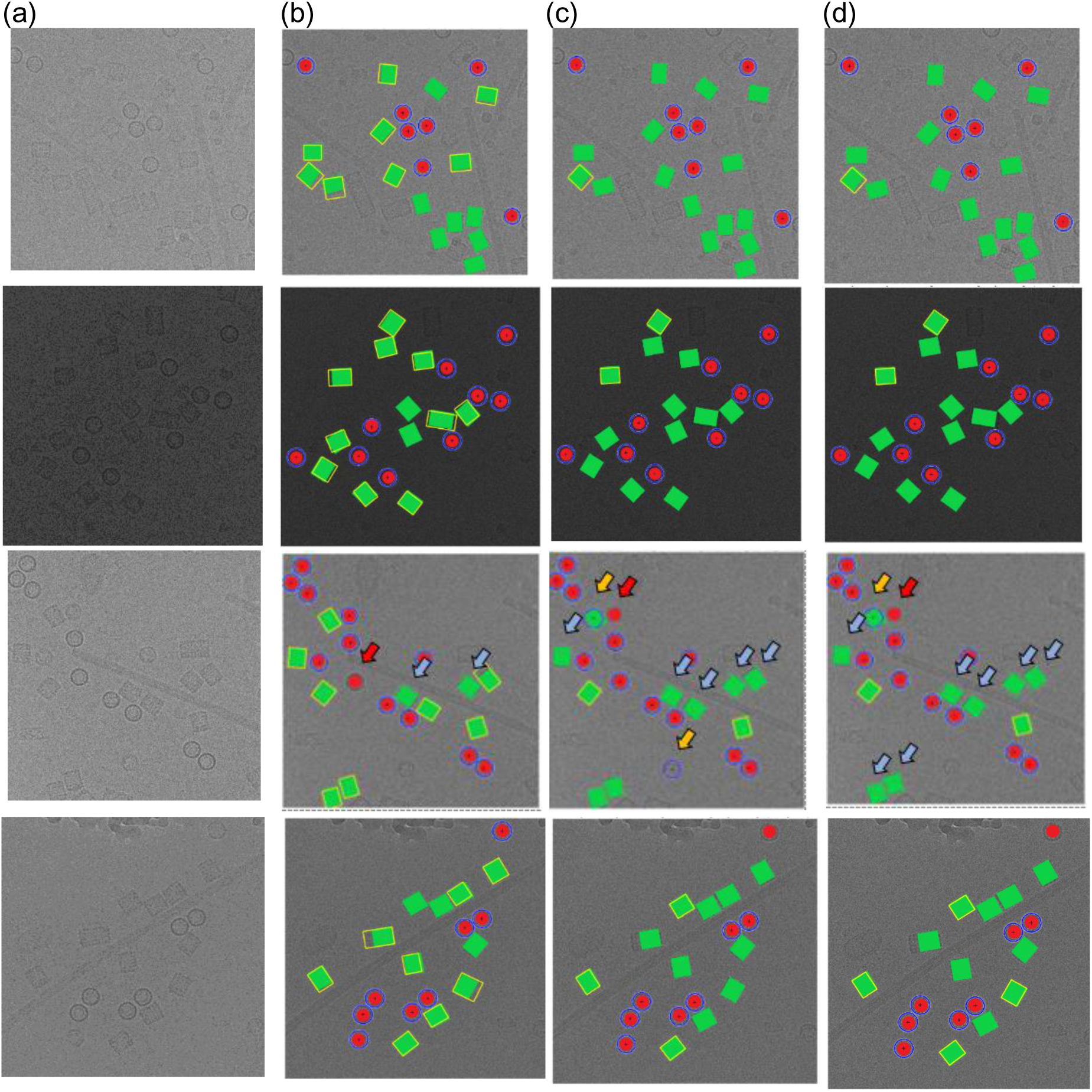
Automated particle picking results for both cases (top and side view) on KLH dataset. (a) The original cryo-EM images form the KLH dataset, (b) Target detection and picking results (top and side particles view) using the ICB clustering algorithm, (c) Target detection and picking results (top and side particles view) using the k-means clustering algorithm, (d) Target detection and picking results (top and side particles view) using the FCM clustering algorithm.

Figure 14(a) shows the original cryo-EM image, while Figure 14(b), (c), and (d) shows the target detection and picking image using ICB, k-means, and FCM respectively. Those examples have been manually labeled in the case of showing the detection and picking performance. The red dots illustrate hand labeling of the circular particles (top view) while the green squares illustrate hand labeling the squares particles (side view), although, the blue circles showing the particle AutoCryoPicker detection and picking results for the top view particles, and the yellow squares showing the side view particles detection and picking results.

Figure 14 illustrates some cases in which AutoCryoPicker failed to detect and pick in both top and side views on the KLH dataset. In the third test example (Figure 14(b), (c), and (d)), there is one top view circular particle not detected by ICB, k-means, and FCM respectively. Figure 14(b) also shows some side view square particles not recognized by ICB clustering. In both cases (top and side view particles), there are almost no false positive detections by ICB clustering, indicating that AutoCryoPicker rarely picked objects from either the background area or icy area. Figure 14(c) and (d) show some side view particles not detected by k-means and FCM respectively. k-means and FCM missed more particles than ICB clustering. They had some false positives (Figure 14(c) and (d)). In one case, a side view was mistakenly detected as a top view, and in another case a background area was detected as a top view.

## Results and discussion

We evaluate the performance of AutoCryoPicker in the three stages according to multiple metrics such as clustering accuracy, particle misclassification (or particles detection) rate, Dice, and time complexity.

### Datasets

Images from two datasets (Apoferritin dataset and Keyhole Limpet Hemocyanin (KLH) dataset) are used to evaluate AutoCryoPicker. The particles in the two datasets are regular shapes, which are ideal for testing AutoCryoPicker because it is designed to detect and pick regular (e.g. circular) particle shapes. Two common shapes of protein particles in cryo-EM images are circles and rectangles.

Apoferritin dataset [34] uses a multi-frame MRC image format (32 Bit Float). The size of each micrograph is 1240 by 1200 pixels. It consists of 20 micrographs each having 50 frames at 2 electrons/A^2/frame, where the beam energy is 300 kV. The particle shape in this dataset is circular.

The Keyhole Limpet Hemocyanin (KLH) dataset from US National Resource for Automated Molecular Microscopy [35] uses a single frame image format in a JPG file format. The size of each micrograph is 2048 by 2048 pixels. It consists of 82 micrographs at 2.2 electrons/A^2/pixel, where the beam energy is 300 120 kV. There are two main types of projection views in this dataset: the top view (circular particle shape) and the side view (square particle shape). The KLH dataset [33] is a standard test dataset for particle picking. The KLH dataset is a challenging dataset because of different specimens (different particles) and confounding artifact (ice contamination, degraded particles, particle aggregates, etc.).

### Evaluation Metrics

In additional to the proposed clustering algorithm (ICB), we select another two main standard cluster algorithms (k-means and FCM). We compare them based on three factors. The first one is the running time. K-means and FCM based pairwise distance comparison is more time consuming. The second one is the accuracy, which includes the clustering accuracy, misclassification rate, dice criteria, precision, recall, and the f1 measure. The third factor is the clustering destabilization. Because K-means and FCM use random selection for cluster initialization, they may group the same points into different clusters in different runs. This requires an extra manual step to select the most appropriate cluster representing particles, which is not fully automated. In contrast, the ICB clustering algorithm is based on computing the interval size to determine the range of the intensity of cluster centers. Therefore, the particles that have the similar intensity values will be grouped into the same cluster.

For the particles clustering stage, we use clustering accuracy and misclassification rate which are defined by Equations (10) and (11), respectively. Each evaluation metric is calculated according to the numbers in a confusion matrix such as the True Positive (TP) which refers to the number of correct detections of positive cases, true Negative (TN) the number of correct detections of negative cases, False Positive (FP) the number of incorrect detections of positive cases and False Negative (FN) the number of incorrect detections of negative cases [38].

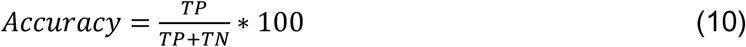

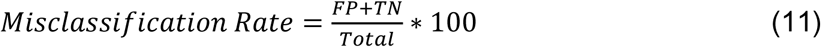

Moreover, Dice Criteria (DIC) is also used for the similarity measure between a cluster image and the Ground Truth (GT). DC is defined by Equation (12) [39]:

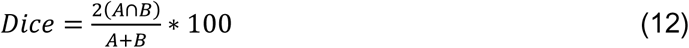

where, *A* is the cluster image and *B* is the ground truth image of *A*. Finally, we use the precision, recall, and F1 measure scores [38] to evaluate the particle picking results in the particle picking stage. The precision, recall, and F measure are defined by Equations (13), (14) and (15), respectively [37]:

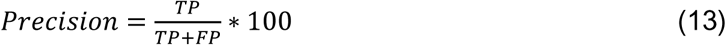

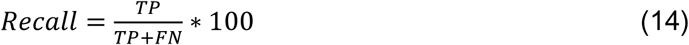

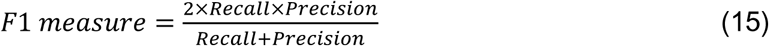

### Particle Clustering, Detection and Picking Results

In order to evaluate the performance of automated particle clustering and picking, we generated a true reference by manually picking the particles on the images. Figure 10(a), and (k) show two different cryo-EM images from the two datasets (Apoferritin and KLH), respectively. The results on one image from the Apoferritin dataset are shown in Figure 10(d), (g), and (j) while the results for KLH dataset are shown in Figure 10(n), (q), and (t). It was demonstrated that most of the particles were correctly picked by AutoCryoPicker. Table 1 reports the recall, precision, accuracy, F1 score, and the running time of AutoCryoPicker based on three clustering algorithms: K-means, FCM, and IBC.

**Table 1.**
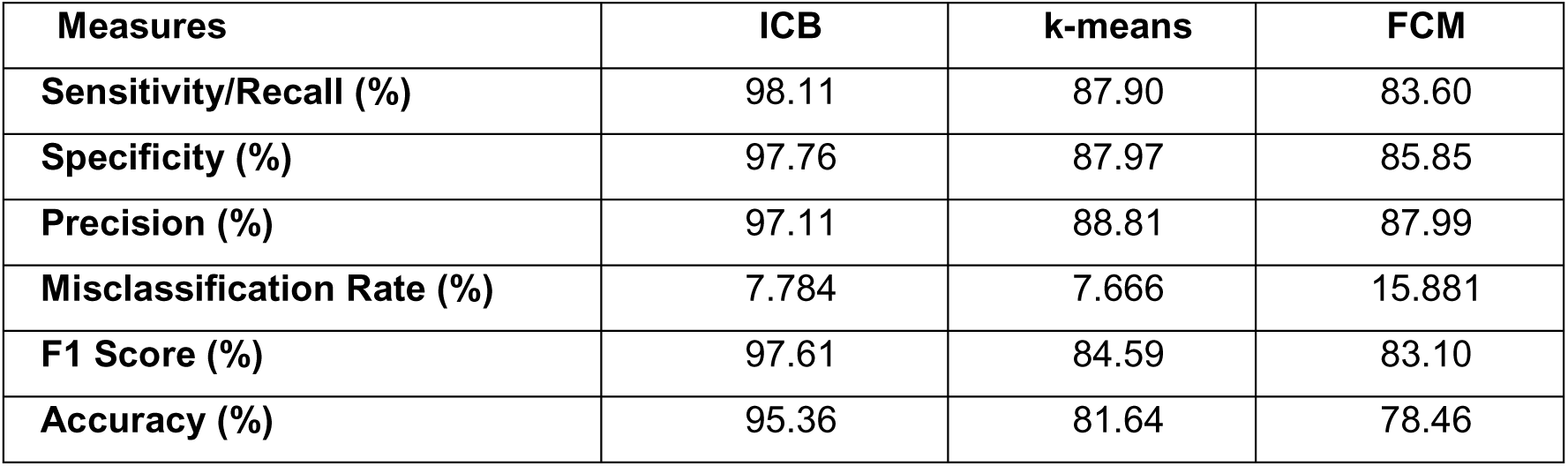
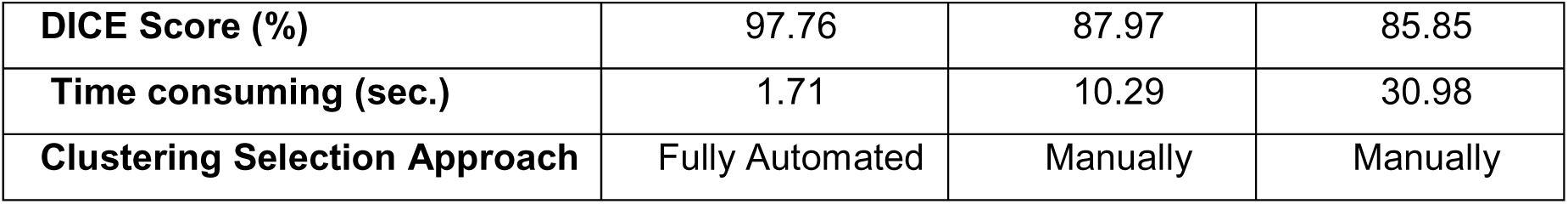
The results of AutoCryoPicker using the three clustering methods on the first dataset (Apoferritin). The table reports the average of the sensitivity or recall, specificity, precision, F1 score, accuracy, DICE score, and the particle clustering time (seconds).

| Measures | ICB | k-means | FCM |
| --- | --- | --- | --- |
| Sensitivity/Recall (%) | 98.11 | 87.90 | 83.60 |
| Specificity (%) | 97.76 | 87.97 | 85.85 |
| Precision (%) | 97.11 | 88.81 | 87.99 |
| Misclassification Rate (%) | 7.784 | 7.666 | 15.881 |
| F1 Score (%) | 97.61 | 84.59 | 83.10 |
| Accuracy (%) | 95.36 | 81.64 | 78.46 |
| **DICE Score (%)** | 97.76 | 87.97 | 85.85 |
| **Time consuming (sec.)** | 1.71 | 10.29 | 30.98 |
| **Clustering Selection Approach** | Fully Automated | Manually | Manually |

On the Apoferritin dataset the AutoCryoPicker based on ICB clustering achieves a higher accuracy of 95.36% than 84.59% and 78.46% of k-means and FCM respectively. Also, ICB ran significantly faster in particles clustering (average time 1.71 seconds versus 10.29 seconds and 30.98 seconds of k-means and FCM, respectively).

Table 2 shows the results on the KLH dataset. AutoCryoPicker based on ICB achieves a higher accuracy 91.82% than that of k-means and FCM (i.e. 87.50% and 80.83% respectively). The average clustering time of the whole dataset using ICB was 4.7 seconds on average, faster than the k-means by 23.8 seconds and 105.8 seconds of the FCM.

**Table 2.** The results of AutoCryoPicker using the three clustering methods on the second dataset (KLH). The table reports the average of the sensitivity or recall, specificity, precision, F1 score, accuracy, DICE score, and the particle clustering time consuming (seconds).

| **Measures** | **ICB** | **k-means** | **FCM** |
| --- | --- | --- | --- |
| **Sensitivity/Recall (%)** | 96.23 | 93.42 | 84.67 |
| **Specificity (%)** | 95.095 | 92.71 | 94.7925 |
| **Precision (%)** | 95.095 | 92.71 | 94.7925 |
| **Misclassification Rate (%)** | 3.77 | 6.58 | 15.33 |
| **F1 Score (%)** | 95.595 | 92.825 | 88.61 |
| **Accuracy (%)** | 91.8275 | 87.5025 | 80.835 |
| **DICE Score (%)** | 95.595 | 92.825 | 89.5 |
| **Time consuming (sec.)** | 4.714643 | 23.8332305 | 105.676302 |

Two different cases from each of the two datasets are illustrated in Figure 15.

Figure 15(a) shows cryo-EM images of a high particle density from the Apoferritin dataset with a low-frequency and Figure 15(b) a cryo-EM image of low SNR. Figure 15(c) and (d) shows two different micrograph cases from the KLH dataset that consist of excessively overlapped particles and some confounding artifact such as ice contamination, degraded particles, and particle aggregates. AutoCryoPicker still performed very well on these cases. Figure 15(e)-(p) show the particle picking results using ICB, k-means, and FCM methods on the two datasets, respectively.

**Figure 15.**
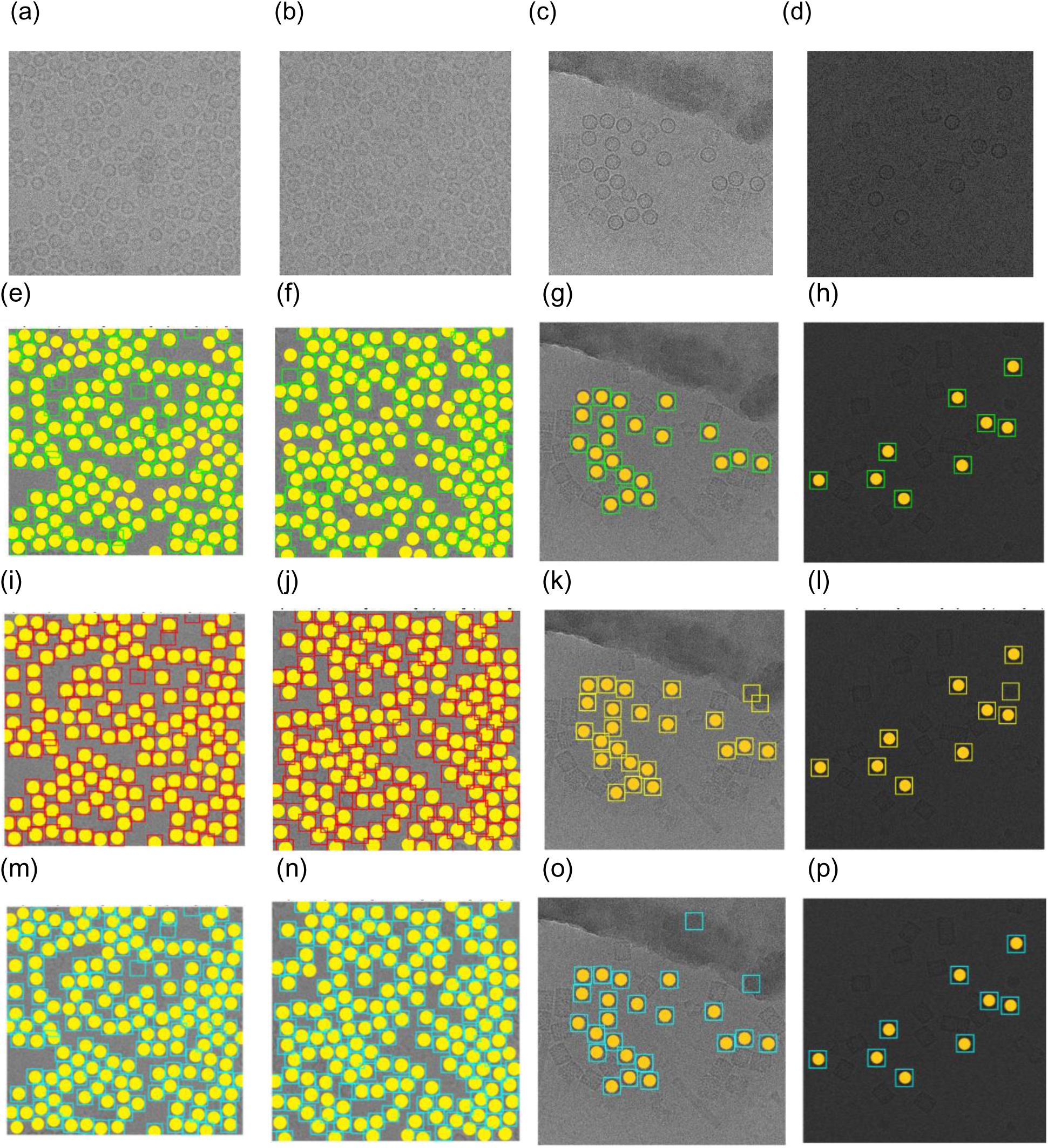
Automated particle picking results on the two datasets. (a) A cryo-EM image with a high identical particle density and a lack low-frequency from the Apoferritin dataset, (b) A low SNR cryo-EM image from the Apoferritin dataset, (c) A micrograph image from the KLH dataset that includes excessively overlapped particles due to confounding artifacts such as ice contamination, degraded particles, and particle aggregates, (d) A micrograph image from the KLH dataset that has a very low spatial density and different intensity levels, and (f) Particle picking results using Intensity Based Clustering Algorithm (ICB) (Apoferritin dataset), and (j) Particle picking results using k-means (Apoferritin dataset), and (n) Particle picking results using FCM (Apoferritin dataset), and (h) Particle picking results using Intensity Based Clustering Algorithm (ICB) (KLH dataset), and (l) Particle picking results using k-means (KLH dataset), and (p) Particle picking results using FCM (KLH dataset).

### Comparison With Another Particle Picking Software

EMAN2 was selected as an example of particle picking software for cryo-EM images [25]. The “e2boxer.py” program of EMAN2 was applied to the same images input to AutoCryoPicker.

For the Apoferritin images, a reference set of 10 particles was selected manually (Figure 16(a), 16(b)) and then automated picking was performed with different threshold values (lower threshold results in more particles picked). For example, use of arbitrarily low threshold values of 0.0 and 0.5 results in most of the valid particles being selected; however, false positives likely corresponding to thick ice were also selected (Figures 16(c), 16(d)). Increasing the threshold to a more reasonable value of 2.3 resulted in no false positives at the expense of leaving several good particles unpicked (Figures 16(e), 16(g)). The lack of particle set completeness is evident by comparison to the ground truth result (Figures 16(f), 16(h)). In comparison, AutoCryoPicker successfully captured all the valid particles on the images without any false positives (Figures 16(m), 16(n)).

**Figure 16.**
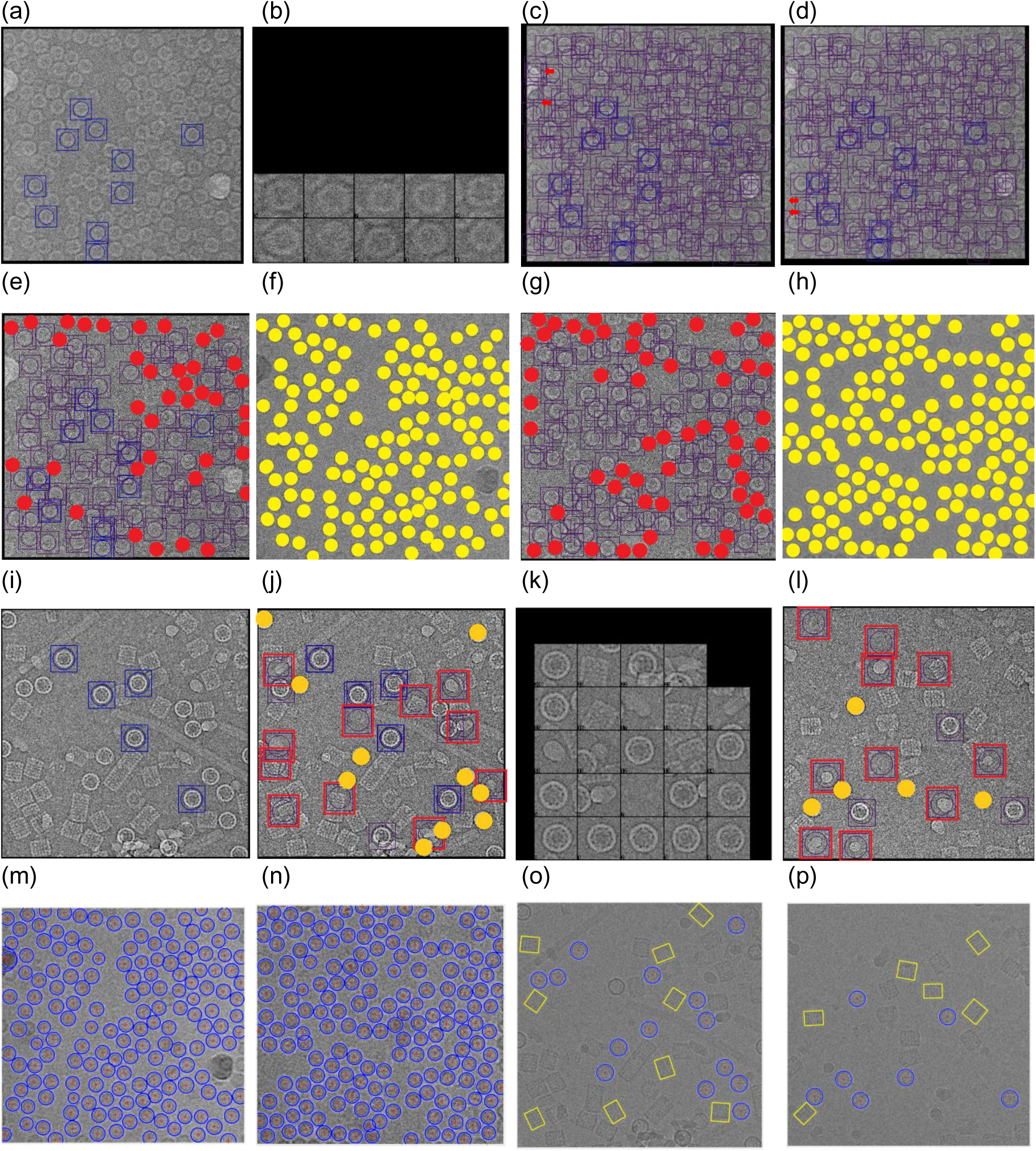
Particle picking using EMAN2. (a) The manually selected reference particles of the Apoferritin dataset that were used for automated particle picking with EMAN2, (b) Zoomed-in view of the reference particles for the Apoferritin dataset, (c) EMAN2 automatic picking result based on threshold value=0.0 using the first tested image of the Apoferritin dataset, (d) EMAN2 automatic picking result based on threshold value=0.5 using the first tested image of the Apoferritin dataset, (e) EMAN2 automatic picking result based the threshold value=2.3 using the first tested image of the Apoferritin dataset. Red dots mark missed particles), (f) Ground truth of first tested image of the Apoferritin dataset. Yellow dots mark valid particles, (g) EMAN2 automatic picking result based the threshold value=2.3 using the second tested image of the Apoferritin dataset. Red dots mark missed particles), (h) Ground truth of second tested image of the Apoferritin dataset. Yellow dots mark valid particles, (i) The manually selected reference particles of the KLH dataset that were used for automated picking of top-view (circular) particles with EMAN2, (j) EMAN2 automatic picking result based the threshold value=0.5 using the first tested image of the KLH dataset. Red squares mark the false positives and the yellow dots the missing particles, (k) Zoomed-in view of the automatically picked particles (threshold value=0.5) for first tested image of the KLH dataset, (l) EMAN2 automatic picking result based the threshold value=0.5 using the second tested image of the KLH dataset. Red squares mark the false positives, and the yellow dots mark the missing particles (top-view), (m) Particle picking result from AutoCryoPicker using the first tested image of the Apoferritin dataset. Red ‘+’ mark the center of each particle and blue circles the top-view detected particles in the cryo-EM image, (n) Particle picking result from AutoCryoPicker using the second tested image of the Apoferritin dataset. Red ‘+’ mark the center of each particle and blue circles the top-view detected particles in the cryo-EM image, (o) Particle picking result from AutoCryoPicker using the first tested image of the KLH dataset. Red ‘+’ marks the center of each particle, blue circles the top-view detected particles in the cryo-EM image, and the yellow squares the side-view detected particles in the cryo-EM image, (p) Particle picking result from AutoCryoPicker using the second tested image from the KLH dataset. Red ‘+’ marks the center of each particle, blue circles the top-view detected particles in the cryo-EM image, and the yellow squares the side-view detected particles in the cryo-EM image.

Similarly, results of using EMAN2 autopicking with the circular particles in the KLH images yielded incomplete recording of the valid particles and several false positives (Figures 16(j), 16 (l)). In contrast, AutoCryoPicker was able to identify almost all of the true particles (both the circular and rectangular projections) in the KLH images, without the generation of false positives (Figures 16(o), 16(p)).

## Conclusions

Accurate particle picking in cryo-EM images still requires substantial human intervention and, therefore, can be labor-intensive and time-consuming. To address this challenge, we develop AutoCryoPicker – a fully automated particle picking approach based on image preprocessing, unsupervised clustering and shape detection. Our experiment shows that the approach can significantly improve signal to noise ratio in cryo-EM images and pick particles rather accurately. Therefore, the automated method can relieve scientists from the laborious work of picking cryo-EM particles and help improve the efficiency and effectiveness of cryo-EM based protein structure determination.

## Abbreviations

Cryo-EM: cryo-electron microscopy
MRC: Medical Research Council
PNG: Portable Network Graphic
Micrograph: Digital image taken through a microscope

## Declarations

### Acknowledgements

Some tests on Apoferritin dataset were carried out by Yuhan Chen.

### Authors Contributions

JC conceived of the project. AA and JC designed the experiment. AA implemented the method and gathered the results. AA, JC, AO and JJT analysed the data. AA and JC wrote the manuscript. All authors edited and approved the manuscript.

### Consent for publication

Not applicable.

### Availability of data and materials

The datasets used in this study and the source code of AutoCryoPicker are available at https://github.com/jianlin-cheng/AutoCryoPicker.

### Ethics approval and consent to participate

Not applicable.

